# Magnesium induces iron starvation and metabolic rewiring to support the viability of cell envelope mutants and antibiotic-stressed cells

**DOI:** 10.64898/2026.08.31.748404

**Authors:** Asher King, Pilar Horigian, Prahathees J. Eswara

## Abstract

Magnesium supplementation permits deletion of otherwise essential genes involved in cell envelope biogenesis in the Gram-positive model bacterium *Bacillus subtilis*. Yet, the specific underlying mechanism has remained elusive. To address this key knowledge gap, we made use of a mutant lacking *ezrA* and *gpsB*. Deletion of both of these genes involved in cell wall synthesis leads to severe growth inhibition which is ameliorated by magnesium addition. Our results indicate that, in the absence of magnesium, this mutant contains elevated levels of labile iron, is impaired in activating the oxidative stress response, and displays extreme sensitivity to iron and manganese intoxication. Intriguingly, we find that an *ezrA* single deletion, but not *gpsB*, exhibits heightened susceptibility to excess iron and manganese. This observation allowed us to investigate the source of toxicity and how EzrA may support metal homeostasis. Our data suggests that the major contributor of ROS is the electron transport system involved in cellular respiration. Both genetic and chemical means to reprogram the cells in favor of fermentation alleviate the metal toxicity in cells lacking *ezrA*. Collectively, our data shows that magnesium limits iron availability and redirects metabolism towards pathways that are preferred during iron scarcity. Consequently, these mechanisms result in reduced ROS production and oxidative stress mitigation. This explains why magnesium supplementation may render essential genes dispensable. In support of this model, we find that addition of magnesium helps cells to circumvent lysis typically caused by the treatment of an antibiotic that disrupts cell wall synthesis. Taken together, our results suggest that unmitigated oxidative stress fueled by labile iron is likely responsible for the detrimental effects of specific gene disruptions and certain antibiotic treatments. By reducing the pool of free iron and reprogramming cellular metabolism, magnesium mitigates oxidative damage and protects cells from ROS-mediated death.

**IMPORTANCE:** Deletion of certain essential genes is possible in *Bacillus subtilis* in the presence of excess magnesium. However, the mechanism behind this is unknown. In this study, we found that cell wall synthesis mutants have increased free reactive iron and are ill-equipped to activate oxidative stress response pathways. Consequently, these mutants are highly susceptible to reactive oxygen species (ROS) specifically stemming from the electron transport system. Our results suggest that magnesium protects cell wall synthesis mutants by limiting free iron availability, activating mechanisms responsible for neutralizing oxidative stress, and redirecting metabolism toward pathways that limit ROS production. Together, these effects alleviate oxidative stress and explain how magnesium helps bypass the requirement of genes that are otherwise considered essential. Finally, our observation that magnesium negatively affects the efficacy of cell wall-targeting antibiotics supports the notion that free iron-dependent ROS serves as a major driver of cell death.

## INTRODUCTION

Divalent cations are vital for cell survival. Chief among them are the redox-active transition metal ions, iron and manganese, which serve as key cofactors in metalloenzymes involved in a variety of fundamental biological processes (Remick & Helmann, 2023). The relatively high reduction potential makes manganese the cofactor of choice in enzymes that respond to oxidative stress (Capek & Vecerek, 2023, Smith *et al*., 2017). In contrast, iron, owing to its low reduction potential, is favored in reactions involving oxygen (Aguirre & Culotta, 2012). As such, iron is almost universally employed in the respiratory electron transport chain (ETC) to generate ATP and support the energy needs of the cell (Kim *et al*., 2012, Goldman *et al*., 2023). However, this process also generates reactive oxygen species (ROS) which can damage DNA, RNA, proteins, and lipids and potentially lead to cell death (Murphy, 2009, Palma *et al*., 2024). Therefore, most organisms encode enzymes such as catalase and superoxide dismutase to neutralize ROS (Imlay, 2013, Xu *et al*., 2025). However, as Fe^2+^ and Mn^2+^ are quite similar, incorrect (or purposeful (Cotruvo & Stubbe, 2012, Imlay, 2014)) incorporation of these metal ions termed mismetallation is known to occur when intracellular metal levels are dysregulated (Keyer *et al*., 1995, Touati *et al*., 1995, Imlay, 2013, Chandrangsu & Helmann, 2016, Smith *et al*., 2017, Fasnacht & Polacek, 2021, Martin & Waters, 2022, Capek & Vecerek, 2023). Thus, if the levels of intracellular metal ions are not finely tuned, it can lead to altered inefficient cellular metabolism and also fuel the generation of ROS. Hence, balancing the divalent cation pool is critical to prevent metal intoxication and achieve the desired cellular response to changing growth conditions. To achieve this, both import and export of metal ions need to be precisely calibrated which is indeed the case in *Bacillus subtilis*, the model Gram-positive bacterium used in this study (Chandrangsu *et al*., 2017). In this organism, manganese toxicity is attributed specifically to defective ETC function (Sachla *et al*., 2021). Intriguingly, the deleterious effect is reversed by increasing the intracellular concentration of another divalent cation, magnesium (Pi *et al*., 2020).

In *B. subtilis*, magnesium supplementation was found to be extremely protective to cells to the extent where certain genes typically considered essential are rendered dispensable (**Fig. 1A**) (Rogers *et al*., 1976, Formstone & Errington, 2005, Leaver & Errington, 2005, D’Elia *et al*., 2006, Schirner & Errington, 2009, Kawai *et al*., 2011, Sassine *et al*., 2020, Tesson *et al*., 2022, Guo & Herman, 2023). Magnesium also supports the growth of certain mutants which otherwise exhibit poor growth or altered cell morphology (Murray *et al*., 1998, Gorke *et al*., 2005, Carballido-Lopez *et al*., 2006, Carballido-Lopez & Formstone, 2007, Matsuoka *et al*., 2011, Suits *et al*., 2026). Notably, all of these genes are directly related to cell envelope biogenesis. Previous studies attributed this beneficial effect of magnesium to inhibition of LytE autolysin (Tesson *et al*., 2022). However, it seems to be rather insufficient as other factors promote LytE to be functional despite the presence of magnesium (Wilson *et al*., 2023). Additionally, cells lacking LytE are also protected by magnesium (Cornilleau *et al*., 2026). Thus, the underlying mechanism that contributes to the protective effects of magnesium supplementation remains to be fully elucidated.

**Figure 1:**
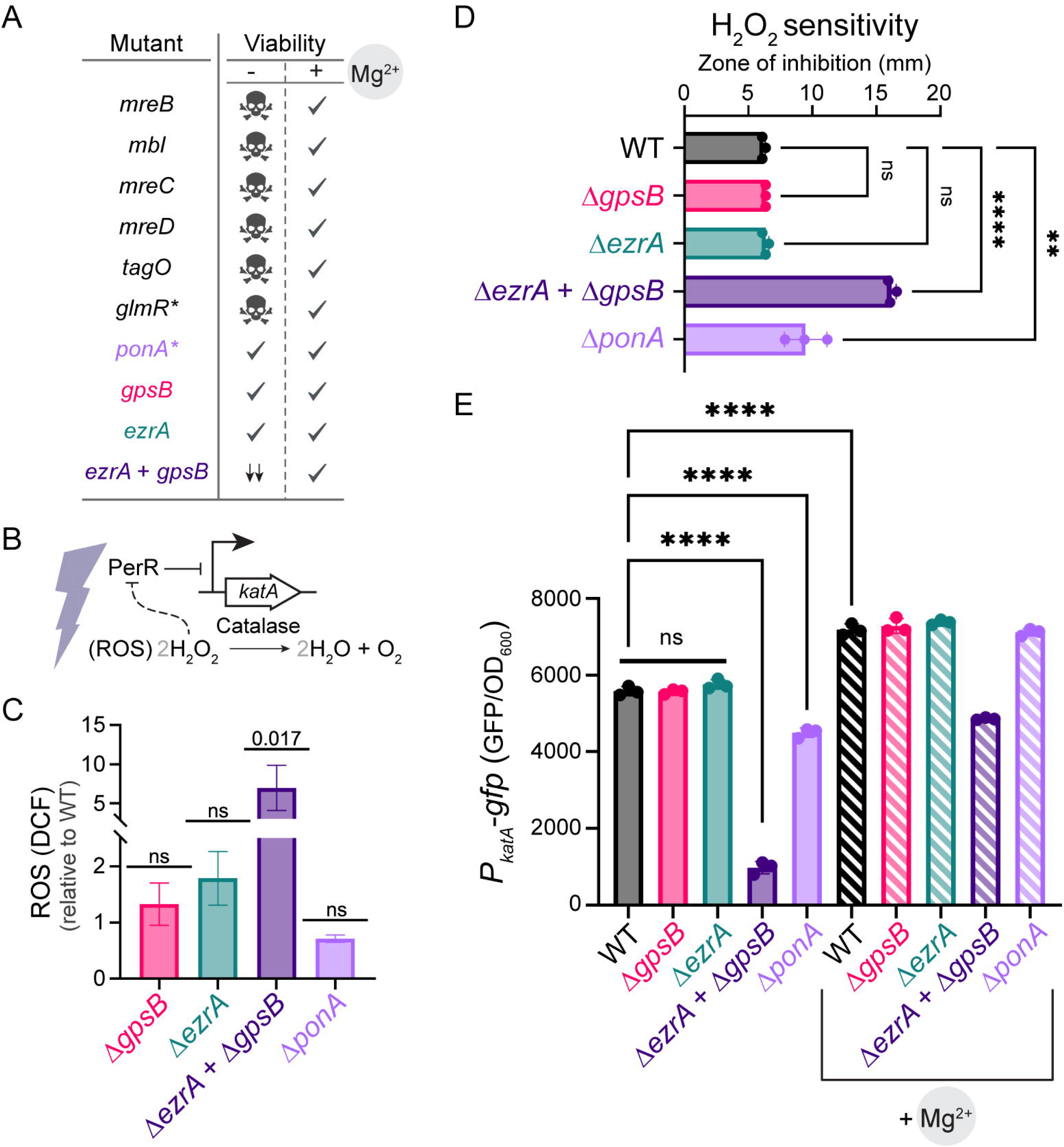
Presence of EzrA and GpsB is required for optimal oxidative stress response. **(A)** Summary of *B. subtilis* cell envelope mutants whose viability or growth depends on magnesium supplementation. Check mark indicates viability, skull symbol denotes loss of viability, and arrows (↓↓) indicate severe growth inhibition. Asterisks denote growth sensitivity only in gluconeogenic conditions and alterations in cell morphology in Δ*glmR* and Δ*ponA* respectively. **(B)** Schematic of PerR-mediated regulation of *katA*. Briefly, iron-bound PerR repression of *katA* is relieved by H_2_O_2_. **(C)** Intracellular ROS levels in Δ*gpsB* (GG13), Δ*ezrA* (AK140), Δ*ezrA* Δ*gpsB* (AK186), and Δ*ponA* (AK286), relative to WT (PY79), were measured using DCFDA assay. Fluorescence value was normalized to OD_600_ and graphed relative to WT (PY79). **(D)** Sensitivity to H_2_O_2_ stress for strains listed in panel C was assessed by standard disc diffusion assay. Zone of inhibition was measured after overnight incubation at 37 °C. **(E)** Activity of the P*_katA_*-*gfp* transcriptional reporter in WT (AK364), Δ*gpsB* (AK356), Δ*ezrA* (AK355), Δ*ezrA* Δ*gpsB* (AK359), and Δ*ponA* (AK371) strains grown in LB medium with or without 25 mM MgSO_4_. Solid bars, no magnesium; hatched bars, magnesium supplemented. GFP fluorescence was normalized to OD_600_. All experiments were performed at least three independent times. The mean and standard deviations are shown. Statistical significance was assessed by one-sample *t*-test (relative to 1) on log_2_-transformed values (C), one-way ANOVA test with Tukey’s post-hoc analysis (D and E). The *p* values are listed (C); or denoted as ** < 0.01, **** < 0.0001, ns - not significant.

Magnesium-mediated protection is also evident in cells harboring deletions of both *ezrA* and *gpsB* (**Fig. 1A**) (Claessen *et al*., 2008). It is known that EzrA and GpsB are interaction partners and that they also mediate interactions with proteins involved in cell division and cell wall synthesis (Haeusser *et al*., 2004, Claessen *et al*., 2008, Tavares *et al*., 2008, Dominguez-Cuevas *et al*., 2013, Bhattacharya *et al*., 2025). Thus, presumably, removal of both *ezrA* and *gpsB* results in dysregulation of multiple processes integral for efficient cell envelope synthesis. As this phenotype is rescued by magnesium supplementation, we utilized this mutant to help uncover the molecular mechanism behind the defensive nature of this divalent cation.

In this report, we reveal that magnesium supplementation leads to increased catalase (*katA*) expression which is normally repressed by PerR (**Fig. 1B**) (Chen *et al*., 1995, Lee & Helmann, 2006). It is established that de-repression of PerR results in a condition that mimics iron starvation due to decreased iron influx and increased iron sequestration (Faulkner *et al*., 2012). Our data suggests that magnesium supplementation restricts respiration, which is conceivable as iron is an important co-factor for the ETC. In the context of an Δ*ezrA* Δ*gpsB* mutant, we observe increased labile iron and defective PerR de-repression as evidenced by decreased *katA* transcription. In addition, we find that this mutant is hypersensitive to iron and manganese intoxication. Further analysis revealed that Δ*ezrA*, but not Δ*gpsB*, exhibits increased sensitivity to excess iron and manganese supplementation. Moreover, we demonstrate that cells devoid of *ezrA* depend on iron (PfeT) and manganese (MneP) efflux to remain viable (Steingard *et al*., 2023). Finally, we show that this severe metal intoxication phenotype can be bypassed with the addition of magnesium or by using other chemical or genetic conditions that limit the use of ROS-producing ETC pathway. Briefly, this was demonstrated by the addition of glucose which facilitates fermentation (Sonenshein, 2007), and by the deletion of *qox* operon (encodes quinol oxidase complex) which restricts respiration (Santana *et al*., 1992, Xu *et al*., 2020, Sachla *et al*., 2021). These observations strongly imply that in cells lacking EzrA and GpsB, and more broadly when cell envelope synthesis is significantly compromised, two intertwined factors lead to ferroptosis-like cell death (Conrad *et al*., 2018, Li *et al*., 2020, Kawai *et al*., 2023, Kwun & Lee, 2023). According to our model, these crucial factors are: (i) elevated labile iron; and (ii) active ETC. Together, this kickstarts a self-amplifying cycle of unmitigated ROS production via Fenton chemistry which eventually leads to cell death (Koppenol, 1993, Imlay, 2013, Capek & Vecerek, 2023).

Based on the evidence presented here, the protective effect of magnesium seems to be due to its ability to limit free iron availability and restrict ROS generative pathways. Next, we hypothesized that (similar to cell wall synthesis mutants), free iron dependent ROS may be the key driver of cell death upon treatment with cell wall targeting antibiotics. We tested this using a beta-lactam antibiotic, cefepime. Astonishingly, in support of our prediction, we see magnesium-treated cells exhibit a substantial increase in viability when compared to wild type cells. Remarkably, we see a similar effect when ETC function is restricted with glucose supplementation or by *qox* deletion. This new understanding may help develop new clinical strategies, as antibiotic recalcitrance may stem from high magnesium or glucose in patients (Lecomte *et al*., 1994, Helaine *et al*., 2024). In sum, we propose that free reactive iron may act as the catalyst of death in cells undergoing respiration with compromised cell envelope and that magnesium protection appears to be mainly mediated by restricting iron availability and disfavoring the ETC pathway.

## RESULTS

### Absence of EzrA and GpsB leads to heightened oxidative stress

As discussed above, the viability of several cell envelope mutants is rescued by magnesium supplementation (**Fig. 1A**). Alternatively, extracellular iron chelation or gene deletions that limit ETC function also support the growth of some of these mutants (Kawai *et al*., 2023). Hence, we argued that magnesium may limit intracellular iron levels and/or reduce endogenous ROS generated by ETC. It is also known that PerR is uniquely responsive to hydrogen peroxide (H_2_O_2_) as it abolishes the repressor function of PerR, specifically when bound to iron (Herbig & Helmann, 2001). This in turn leads to the upregulation of *katA* which encodes for catalase which is responsible for detoxifying H_2_O_2_ (**Fig. 1B**). Additionally, catalase neutralizes the threat posed by H_2_O_2_ in initiating Fenton chemistry (Haber & Weiss, 1934). Based on this, we postulated that cells lacking both *gpsB* and *ezrA* may accumulate ROS to a higher degree. To assess this, we utilized the ROS-sensitive reagent 2’,7’-dichlorodihydrofluorescein (DCF) diacetate, which upon oxidization becomes fluorescent (Sachla *et al*., 2024). We note the DCF fluorescence, normalized to WT, was significantly elevated in the absence of EzrA and GpsB (**Fig. 1C**). However, individual deletions did not result in an appreciable increase in the intracellular ROS levels. Nonetheless, this result supports our hypothesis by revealing that the Δ*ezrA* Δ*gpsB* double-deletion strain has elevated levels of reactive oxygen radicals.

Given that an *ezrA* and *gpsB* double-deletion mutant intrinsically has high ROS, we suspected that this mutant may be hypersusceptible to oxidative stress. To test this, we conducted a disc-diffusion assay with H_2_O_2_ (**Fig. 1D**). The class A penicillin binding protein PBP1 (encoded by *ponA*) is a direct interaction partner of both EzrA and GpsB (Claessen *et al*., 2008), and cells lacking *ponA* undergo magnesium-dependent cell shape correction (Murray *et al*., 1998). Hence, we also included a *ponA* deletion mutant in our experiments. Our results reveal that mutants lacking *ezrA* or *gpsB* resemble our wild type (WT) control, while deletion of both *ezrA* and *gpsB* resulted in increased H_2_O_2_ sensitivity. Intriguingly, cells lacking *ponA* also displayed a modest but significant increase in sensitivity. Nonetheless, as predicted, cells lacking *ezrA* and *gpsB* are indeed extremely sensitive to oxidative stress.

As PerR is the main transcription factor responsible for responding to H_2_O_2_ (**Fig. 1B**) (Herbig & Helmann, 2001), we utilized a *P_katA_-gfp* transcriptional reporter (Hoover *et al*., 2010), to investigate PerR function in our mutant strain backgrounds. Consistent with previous observations (Herbig & Helmann, 2001, Hoover *et al*., 2010), *katA* expression is seen in WT and a similar level of transcriptional activity is recorded in the Δ*ezrA* and Δ*gpsB* mutant backgrounds (**Fig. 1E**). Strikingly, the promoter activity is severely dampened in cells lacking *ezrA* and *gpsB*, while it is only modestly reduced in cells lacking *ponA*. This indicates that the repressor function of PerR is strengthened in the Δ*ezrA* Δ*gpsB* double-deletion mutant. Given that magnesium alleviates the growth impairment of this mutant (Claessen *et al*., 2008), we wondered whether magnesium increases *katA* expression in this strain. To our surprise, our results reveal that this is indeed the case not only for the Δ*ezrA* Δ*gpsB* strain but also for the rest of the strains including WT where we see a significant increase in *P_katA_-gfp* activity upon magnesium supplementation. As excess iron or manganese typically repress this promoter (**Fig. S1D**) (Herbig & Helmann, 2001), we infer that both metal ions are unavailable within the cell to facilitate PerR repression. Taken together, our results suggest that deletion of both *ezrA* and *gpsB* leads to H_2_O_2_ hypersensitivity presumably due to impaired PerR function and that magnesium corrects this defect.

### Disruption of cell wall synthesis results in increased availability of labile iron

It is known that H_2_O_2_-induced PerR de-repression of *katA* occurs when it is bound to iron, but not manganese (Herbig & Helmann, 2001). As ferroproteins sequester iron intracellularly, free iron is usually limited in the cell which in turn prevents its participation in Fenton chemistry (Andrews *et al*., 2003, Moore & Helmann, 2005). As we noticed strong repression of *P_katA_-gfp* (**Fig. 1E**) in the Δ*ezrA* Δ*gpsB* strain, we presumed this might be due to excess iron in the cell. Therefore, we measured the total intracellular iron content using Ferene-S assay (Chung, 1985, Mashruwala *et al*., 2015). This revealed that cells lacking both *ezrA* and *gpsB* have increased iron levels compared to WT and corresponding single mutants (**Fig. 2A** and **Fig. S1A**). Intriguingly, a *ponA* deletion strain also displayed a slightly elevated amount of total iron but not to the same extent as the Δ*ezrA* Δ*gpsB* double mutant. To further distinguish whether the intracellular iron is present in an unbound form or safely sequestered by ferroproteins, we made use of the antibiotic streptonigrin (SN). It has been established that SN specifically requires free iron for its toxic effects (Yeowell & White, 1982, Mashruwala *et al*., 2015, Guan *et al*., 2015, Gupta & Imlay, 2023). Briefly, increased levels of unbound iron would react with SN and result in heightened toxicity (**Fig. 2B**). To investigate this, we spotted the 10-fold serial dilutions of WT or mutant strains on a plate containing 0, 17.5, 50, and 60 ng/ml of SN. Our experiments revealed that in comparison to the WT, the Δ*ezrA* Δ*gpsB* strain is extremely sensitive to SN even at 17.5 ng/ml. This indicates that cells lacking both *ezrA* and *gpsB* have elevated levels of free reactive iron. We also find that Δ*ezrA* and Δ*ponA* single-deletion mutants exhibit heightened sensitivity that becomes more obvious at 50 ng/ml. This sensitivity is also recapitulated in liquid culture (**Fig. S1B**). In contrast, the sensitivity of Δ*gpsB* cells resembled that of WT and their growth was impaired only at the highest concentration of SN tested (60 ng/ml). Intriguingly, although the total iron level is comparable to WT for *ezrA* and *ponA* single mutants (**Fig. 2A**), their heightened sensitivity to SN implies that iron in those strains is likely present in its unbound labile form. Importantly, SN sensitivity was reverted to WT range in Δ*ezrA* cells through complementation (**Fig. S1C**). This indicates a specific role for EzrA in regulating the labile iron pool. Together, the data shown in **Fig. 2C** reveals that the intracellular labile iron level is elevated in the absence of EzrA or PBP1.

**Figure 2:**
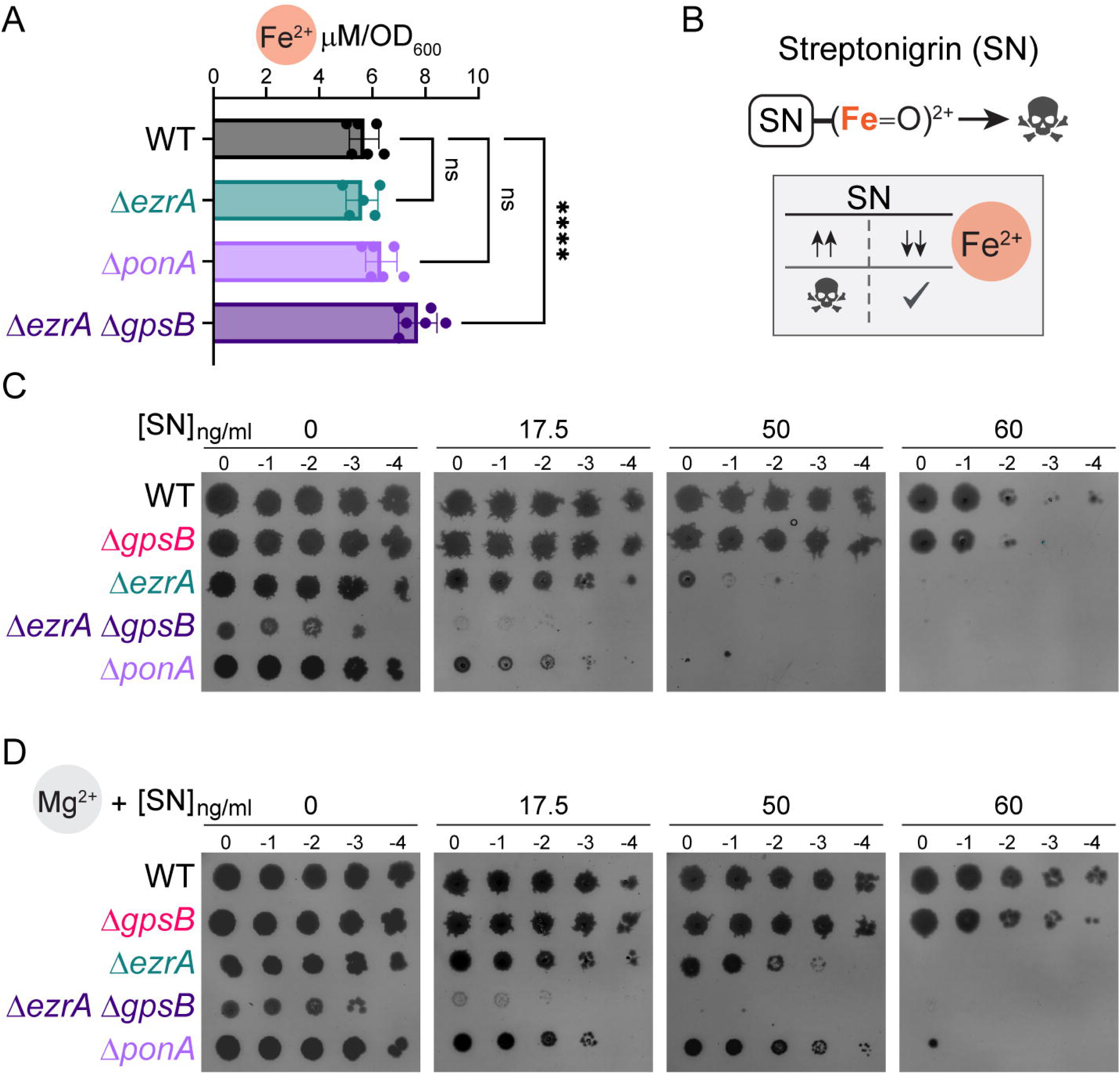
Free reactive iron level is moderately elevated in *ezrA* and *ponA* single-deletion mutants and highly elevated in *ezrA gpsB* double-deletion strain. **(A)** Total intracellular iron (bound and unbound iron) content of WT (PY79), Δ*ezrA* (AK140), Δ*ezrA* Δ*gpsB* (AK186), and Δ*ponA* (AK286) were determined using Ferene-S assay. The mean and standard deviations of 3 biological replicates are indicated. Statistical significance was calculated by one-way ANOVA test with Tukey’s post-hoc analysis; \*\*\*\**p* < 0.0001; ns, not significant. Data corresponding to Δ*gpsB* is shown in Fig. S1A. **(B)** Streptonigrin (SN) toxicity depends on free iron. Elevated levels (↑↑) of labile iron facilitate SN-mediated killing through the formation of highly-reactive ferryl radical. In contrast, limited availability (↓↓) of unbound iron allows cells to be viable. **(C)** Assay for streptonigrin sensitivity. Ten-fold serial dilutions (10^0^-10^-4^) of WT (PY79), Δ*gpsB* (GG13), Δ*ezrA* (AK140), Δ*ezrA* Δ*gpsB* (AK186), and Δ*ponA* (AK286) were spotted on LB medium containing 0, 17.5, 50, or 60 ng/ml SN. **(D)** As in (C), except plates were additionally supplemented with 25 mM MgSO_4_. Representative plate images (C and D) of at least 3 independent experiments are shown.

PerR de-repression is known to result in iron stringency (Faulkner *et al*., 2012). This happens in the presence of magnesium supplementation as evidenced by increased *katA* promoter activity (**Fig. 1E**). Therefore, we hypothesized that magnesium supplementation may alleviate SN toxicity by inducing iron starvation. As revealed in **Fig. 2D**, this is in fact the case for all strains tested. Specifically, the growth impairment of WT and Δ*gpsB* seen at 60 ng/ml SN was abrogated in the presence of magnesium. Similarly, we observe significant growth improvement for Δ*ezrA* and Δ*ponA* strains (compare 50 ng/ml plates in **Fig. 2C** and **2D**). Nonetheless, at 60 ng/ml SN, magnesium supplementation is insufficient to restore the viability of these mutants. In contrast, for the Δ*ezrA* Δ*gpsB* double knockout strain, we only see a subtle improvement in growth at 17.5 ng/ml. This suggests that magnesium supplementation is inadequate to protect cells against SN when neither EzrA nor GpsB are present. Collectively, our data supports the notion that magnesium supplementation reduces the availability of free iron and protects the cells against the toxic effects of SN.

### EzrA helps mitigate metal-induced toxicity

As our experiments with SN hinted at a potential dysregulation of iron homeostasis in our mutants, we postulated that they may not be able to cope with metal intoxication (Andrews *et al*., 2003, Steingard *et al*., 2023). Importantly, if divalent cation homeostasis is impaired in our mutants, then excess labile iron would exacerbate ROS production via Fenton chemistry and alternatively excess manganese would inhibit oxidative stress response pathways (Herbig & Helmann, 2001). As such, the viability of the mutants would be hampered in either condition. To test this, we grew WT and mutant strains on plates supplemented with either 2 mM ferric chloride or 0.4 mM manganese chloride (**Fig. 3A**). As expected, both WT and Δ*gpsB* strains maintain viability on both plates without any obvious growth defects. However, surprisingly, we find cells lacking *ponA* display no apparent signs of growth inhibition in the presence of iron or manganese supplementation. This suggests that although the level of labile iron is relatively high in Δ*ponA* cells (**Fig. 2C**), they retain the mechanisms needed to circumvent metal intoxication. Conversely, we see that colonies of Δ*ezrA* mutant are translucent after a 24-h incubation period, indicating growth impediment specifically in the presence of excess iron or manganese. However, growth of this mutant appears to eventually recover after 42 h. This implies that the metal intoxication response is perhaps delayed in the absence of EzrA. In striking contrast, Δ*ezrA* Δ*gpsB* mutants are unable to recover even after longer incubation. Hence, GpsB, while normally dispensable, becomes important when EzrA is absent to help counter metal toxicity.

**Figure 3:**
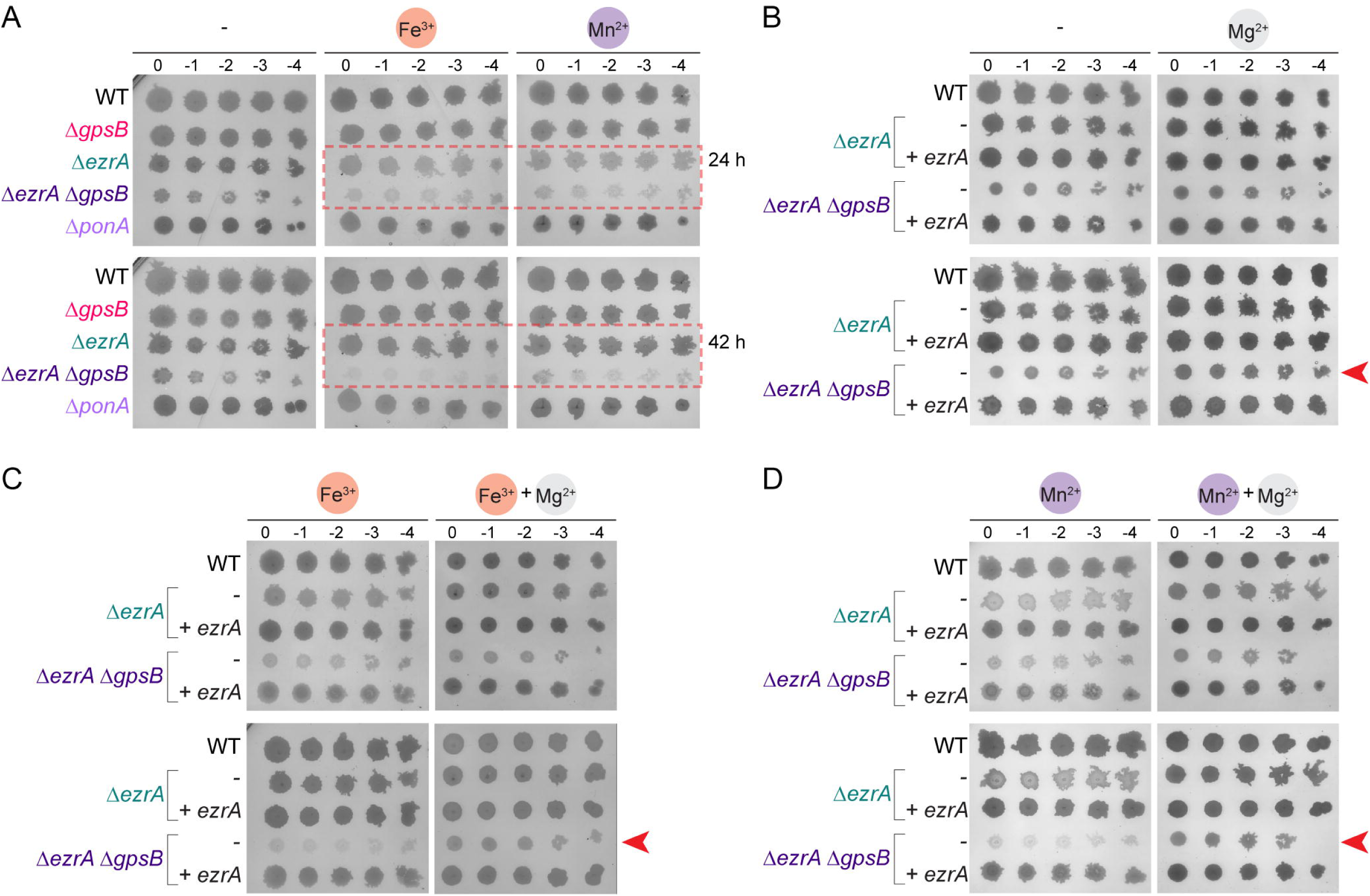
Magnesium alleviates metal-induced toxicity in *ezrA* and *ezrA gpsB* mutants. **(A)** Spot titer assay of WT (PY79), Δ*gpsB* (GG13), Δ*ezrA* (AK140), Δ*ezrA* Δ*gpsB* (AK186), and Δ*ponA* (AK286) strains grown on LB media alone or supplemented with 2 mM FeCl_3_ or 0.4 mM MnCl_2_. Red dashed boxes highlight the inhibited growth of Δ*ezrA* and the persistent growth defect of Δ*ezrA* Δ*gpsB*. **(B**) As in (A), growth of WT (PY79), Δ*ezrA* (AK140), Δ*ezrA ezrA^+^* (AK477), Δ*ezrA* Δ*gpsB* (AK186), and Δ*ezrA* Δ*gpsB ezrA^+^* (AK478) strains on LB media with or without 25 mM MgSO_4_. **(C)** Strains listed in (B), grown on plates containing 2 mM FeCl_3_ with or without 25 mM MgSO_4_. **(D)** Similarly strains listed on (B), grown on plates containing 0.4 mM MnCl_2_ with or without 25 mM MgSO_4_. Plate pictures are representative of at least 3 independent experiments imaged after 24 h (top panels) and 42 h (bottom panels) incubation. Arrowheads indicate growth recovery in the presence of magnesium.

As magnesium supplementation has been shown to reduce manganese sensitivity (Sachla *et al*., 2024), we explored this possibility in the context of our cell wall mutants. As we see magnesium promotes PerR de-repression presumably by limiting iron and manganese availability in cells (**Fig. S1D**), it may equip Δ*ezrA* and Δ*ezrA* Δ*gpsB* mutant strains with tools needed to address metal toxicity. To assess this, we repeated the experiment with iron and manganese both in the absence and presence of magnesium. As previously reported (Claessen *et al*., 2008), the growth impairment of the Δ*ezrA* Δ*gpsB* strain is alleviated by magnesium supplementation as evidenced by increased opacity in the presence of this divalent cation or by *ezrA* complementation (**Fig. 3B**). Curiously, we see that the pale colony phenotype of the Δ*ezrA* mutant seen in the presence of excess iron (**Fig. 3C**) or manganese (**Fig. 3D**) was essentially reversed by the addition of magnesium even after a shorter 24 h incubation. Similarly, for cells harboring an *ezrA* and *gpsB* double-deletion, the lack of growth recovery we observe in the presence of iron or manganese was abrogated by magnesium supplementation and is apparent at both 24 h and 42 h (**Fig. 3BCD**; see arrowheads). Again, this hypersensitivity to metal ions is also reversed by re-introduction of *ezrA* alone suggesting that EzrA is a major contributor to metal homeostasis. Overall, this set of results further confirms that magnesium negatively affects the intracellular availability and/or the effects of toxic metal ions.

### Efflux of metal ions is crucial for survival in cell wall synthesis mutants

As the protective effect of magnesium appears to stem from safe iron management, we hypothesized that the function of one or more iron efflux pumps is impaired when cell wall synthesis is compromised. This in turn leads to the deleterious phenotypes we observe. It has already been demonstrated that PfeT (also regulated by PerR) is the main exporter of iron while the manganese exporters MneP and to a lesser extent MneS may also be utilized to maintain the proper iron to manganese ratio (Steingard *et al*., 2023). Therefore, based on SN sensitivity and metal intoxication (**Figs. 2C** and **3A**), we chose *ezrA* and *ponA* null mutants for further scrutiny. Briefly, we introduced additional deletion of *pfeT*, *mneP*, or *mneS* in Δ*ezrA* or Δ*ponA* strain backgrounds and tested their sensitivity to increasing concentrations of iron (FeCl_3_) and manganese (MnCl_2_) after a 42-h incubation.

At the highest iron concentration (4 mM), WT is able to form colonies at all dilutions while the growth of cells harboring *pfeT* deletion is severely impaired (**Fig. 4A**), as expected (Guan *et al*., 2015, Steingard *et al*., 2023). Conversely, at this concentration (4 mM) we see translucent colonies for the Δ*ezrA* strain, implying poor growth recovery even after 42 h. Remarkably, we find that the Δ*pfeT* Δ*ezrA* mutant is highly sensitive to iron intoxication even at the lowest concentration tested (see arrowheads i). This observation suggests that when EzrA is absent cells rely on PfeT to export iron to maintain viability. Similarly, while *ponA* null cells are quite tolerant to excess iron, the growth of the Δ*pfeT* Δ*ponA* mutant is highly affected (see arrowheads ii). However, heightened sensitivity to iron was not seen in mutants lacking both *gpsB* and *pfeT* (**Fig. S2A**, top panel). Thus, our data reveal that cells lacking *ezrA* or *ponA*, but not *gpsB*, highly depend on PfeT for iron efflux to avoid cell death.

**Figure 4:**
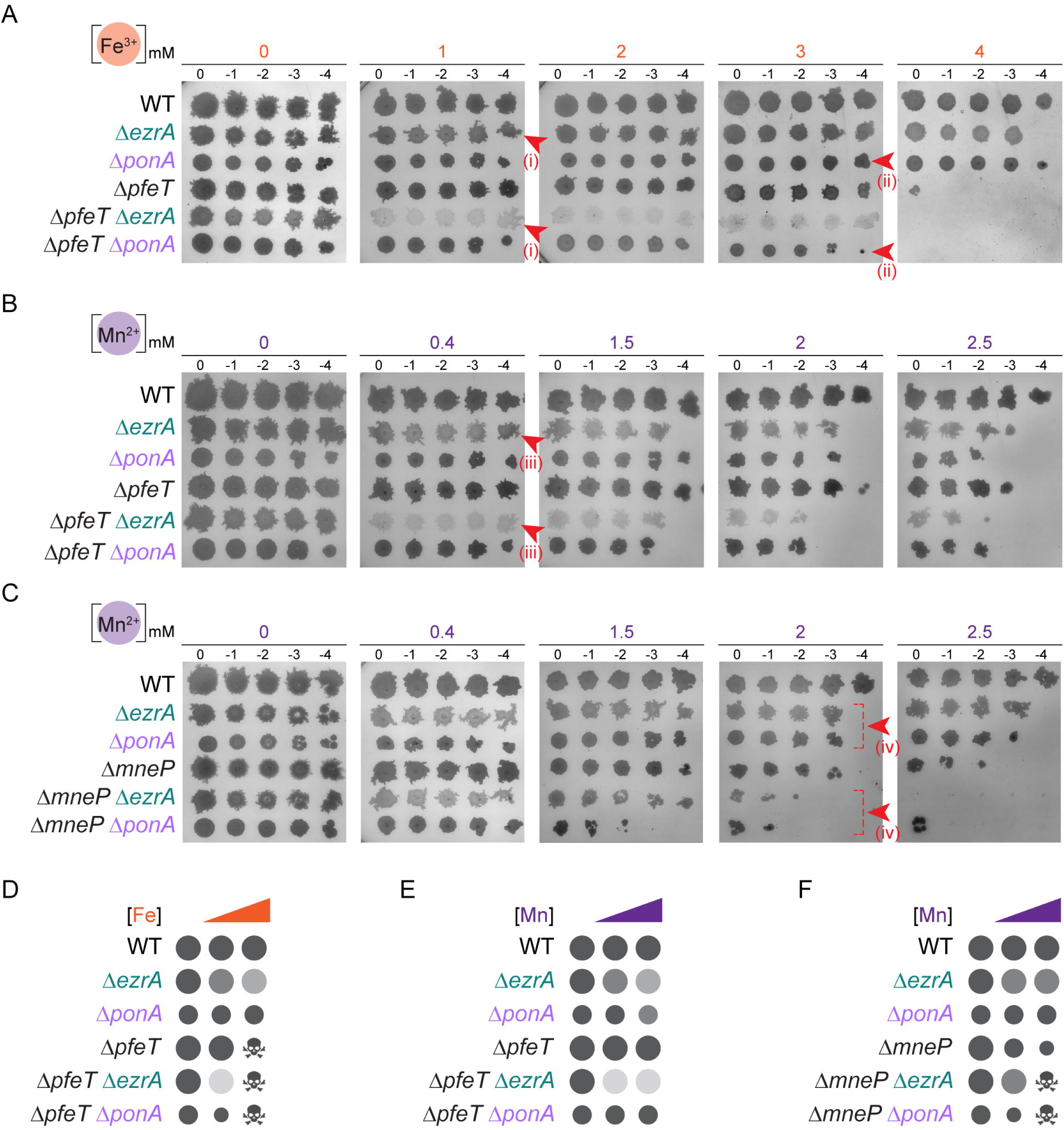
Metal homeostasis is impaired in mutants lacking *ezrA* or *ponA* and metal intoxication is worsened upon the removal of metal efflux pumps in these strains. **(A)** Spot titer assay of WT (PY79), Δ*ezrA* (AK140), Δ*ponA* (AK286), Δ*pfeT* (AK501), Δ*pfeT* Δ*ezrA* (AK504) and Δ*pfeT* Δ*ponA* (AK510) strains grown on LB medium containing 0, 1, 2, 3, or 4 mM FeCl_3_ and imaged after 42 h. Red arrowheads (i) and (ii) mark the iron hypersensitivity of Δ*pfeT* Δ*ezrA* (1 mM) and Δ*pfeT* Δ*ponA* (3 mM) respectively. **(B)** The strains listed in panel (A), were grown on plates containing 0, 0.4, 1.5, 2, or 2.5 mM MnCl_2_. Arrowheads (iii) mark the enhanced manganese sensitivity of Δ*pfeT* Δ*ezrA* at 0.4 mM. **(C)** As in (B), with WT (PY79), Δ*ezrA* (AK140), Δ*ponA* (AK286), Δ*mneP* (AK498), Δ*mneP* Δ*ezrA* (AK513), and Δ*mneP* Δ*ponA* (AK532) strains. Red arrowheads (iv) point to the growth impairment, due to deletion of *mneP*, in Δ*ezrA* and Δ*ponA* mutants grown at 2 mM Mn^2+^. **(D-F)** Schematic summaries of the data in (A-C), respectively. Circle size and translucency represent relative colony size and changes in metal sensitivity across increasing metal concentration; skull symbols denote loss of viability. All images are representative of at least three independent experiments.

As manganese is known to repress *pfeT* expression (Pinochet-Barros & Helmann, 2020), we assessed the sensitivity of the mutants generated to this divalent cation (**Fig. 4B**). WT growth is unaffected even at the highest concentration tested (2.5 mM). Deletion of *pfeT* also appears to not affect growth significantly in the presence of excess manganese (Guan *et al*., 2015). In contrast, we observe severe (translucent colonies) and moderate growth inhibition for Δ*ezrA* and Δ*ponA* strains respectively. Surprisingly, additional deletion of *pfeT* further hampers the growth of the Δ*ezrA* mutant (see arrowheads iii), but not cells devoid of *ponA*. This implies that cells lacking *ponA* may utilize an alternate pathway to circumvent manganese intoxication. We observe that cells harboring deletion of both *gpsB* and *pfeT* do not exhibit manganese sensitivity (**Fig. S2A**, bottom panel). This set of data reveals that the function of PfeT in mitigating manganese toxicity becomes paramount in the absence of EzrA.

Next, we tested the roles of MneP and MneS in managing manganese or iron intoxication in our mutant backgrounds. Our results show that *mneP* or *mneS* deletion individually or in combination with *ezrA*, *ponA*, or *gpsB* does not cause any apparent growth inhibition in the presence of excess iron (**Fig. S2BCD**). In contrast, when grown in the presence of excess manganese, we observe moderate growth inhibition for both Δ*mneP* Δ*ezrA* and Δ*mneP* Δ*ponA* (**Fig. 4C**; see arrowheads iv). However, these strains experience near-complete growth inhibition at 2.5 mM MnCl_2_. We also see a more modest growth impairment at higher concentrations of manganese for Δ*mneS* Δ*ezrA* and Δ*mneS* Δ*ponA* (**Fig. S2D**, compare rows marked by arrowheads ii and iii). We also investigated manganese sensitivity in mutants lacking *gpsB*. While the growth profile of the *mneP gpsB* double-deletion strain resembled that of the Δ*mneP* mutant (**Fig. S2C**), we do see that cells lacking *gpsB* become more sensitive to manganese in the absence of MneS (**Fig. S2C**, see arrowheads i). Together, our data suggests that severe manganese toxicity in *ezrA* and *ponA* null mutants is alleviated by MneP activity.

We then tested whether the protective effect of magnesium seen for cells lacking *ezrA* against iron and manganese intoxication (**Fig. 3**) would also protect the hypersensitivity of the Δ*pfeT* Δ*ezrA* mutant. Notably, magnesium supplementation was able to enhance the growth of Δ*pfeT* Δ*ezrA* strain on both iron and manganese (**Fig. S3AB**). Additionally, we see similar magnesium protection for the *mneP ponA* double-deletion mutant grown on manganese (**Fig. S3C**). Collectively, our results illustrate that the presence of excess iron or manganese inhibits the growth of cells lacking *ezrA* and *ponA*. This growth impairment is exacerbated with additional deletion of *pfeT*, especially in cells lacking *ezrA* upon exposure to excess iron or manganese (**Fig. 4DE**). It appears that MneP and MneS are not needed to manage iron intoxication in our genetic backgrounds, presumably due to the presence of PfeT. Nevertheless, we see that MneP is the major player in preventing manganese intoxication in *ezrA* and *ponA* deletion mutants (**Fig. 4F**). We also note that MneS plays a role, albeit a minor one, in addressing manganese toxicity in our mutant strains. Again, the toxic effects of manganese seen in the Δ*mneP* Δ*ezrA* and Δ*mneP* Δ*ponA* mutants are also masked by magnesium. Together with results discussed previously, it appears that magnesium may limit the availability of iron and manganese.

### ETC function renders Δ*ezrA* cells susceptible to metal induced toxicity

*B. subtilis* is capable of harnessing both oxidative phosphorylation (ETC pathway; shown in pink in **Fig. 5A**) and overflow metabolism (fermentation; shown in yellow) pathways to produce energy, manage oxidative stress, and remain viable (Sonenshein, 2007). It was previously shown that manganese intoxication is due to dysregulated ETC function and that this toxicity can be ameliorated by the removal of the terminal menaquinol oxidase encoded by the *qoxABCD* operon (Sachla *et al*., 2021). Deletion of the *qox* operon also protects against zinc toxicity which is mediated by excess heme iron produced due to PerR dysfunction (Chandrangsu & Helmann, 2016, Sachla *et al*., 2021). As we see severe iron and manganese sensitivity in the absence of EzrA (**Fig. 2C** and **Fig. 3A**), we hypothesized that deletion of the *qox* operon would also protect Δ*ezrA* cells from metal intoxication. As shown in **Fig. 5B**, this is indeed the case for both iron and manganese. This suggests that in the absence of EzrA, the Qox complex is the main source of iron and manganese toxicity.

**Figure 5:**
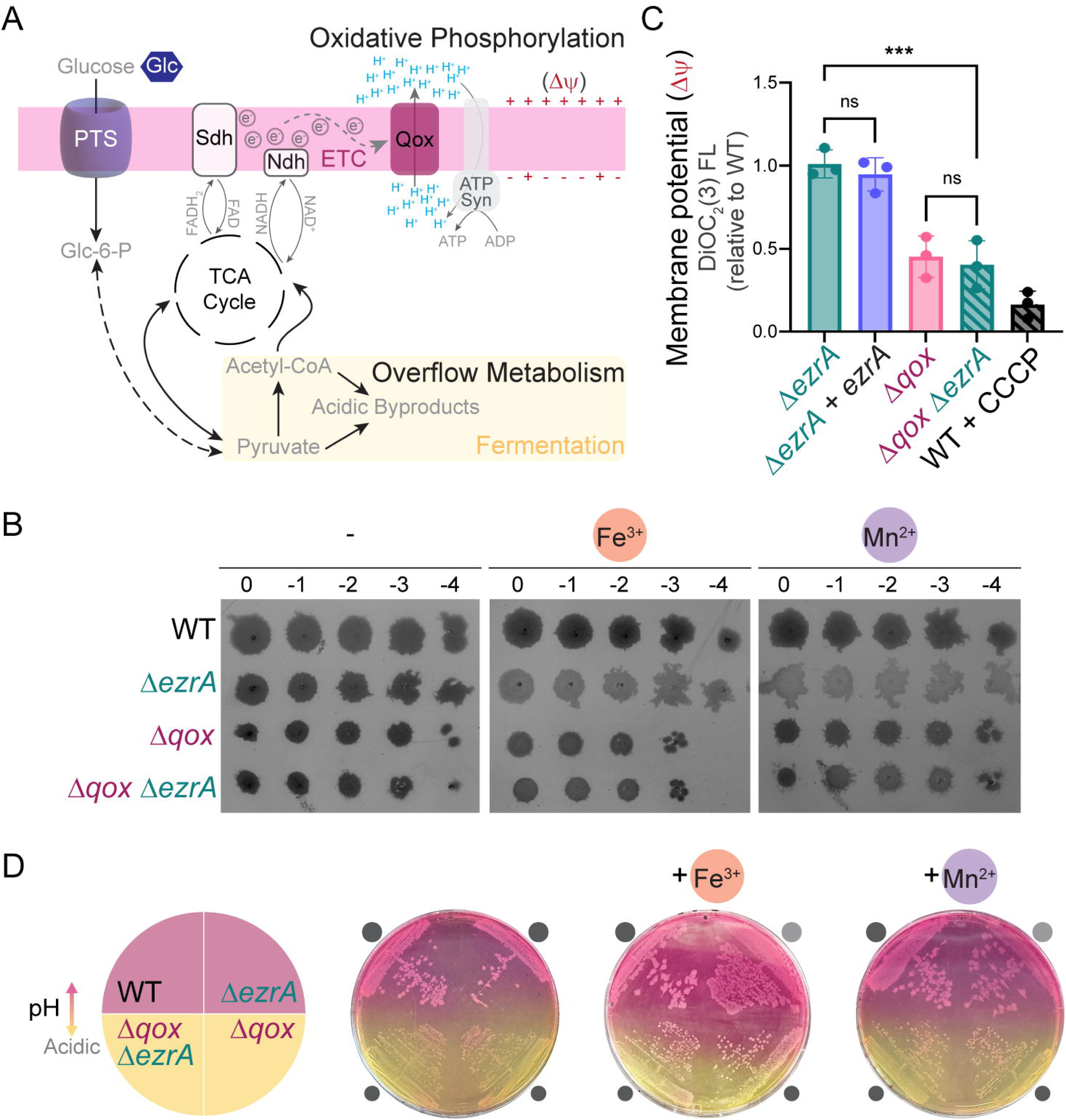
Removal of quinol oxidase complex eases metal-induced toxicity in cells lacking *ezrA*. **(A)** Schematic of the *B. subtilis* energy generative pathways and proteins involved that are relevant to this study: oxidative phosphorylation (respiratory metabolism) and overflow metabolism (fermentation). Glucose (Glc) is imported via the PTS complex and enters glycolysis. High-energy electron carriers such as NADH and FADH_2_ produced during glycolysis and tricarboxylic acid (TCA) cycle are used to fuel the electron transport chain (ETC) via NADH dehydrogenase (Ndh) and other proteins/complexes such as succinate dehydrogenase (Sdh) respectively. These electrons are then passed to the terminal quinol oxidase (Qox) complex which reduces oxygen (terminal electron acceptor) to water and couples this energy to pump protons (H^+^) to generate and maintain proton gradient (ΔpH) across the cell membrane. This process together with other ion channels, for example corresponding to potassium, work together to generate the membrane potential (Δψ). Thus, the proton motive force (PMF) which powers the ATP synthase machinery to produce ATP integrates both ΔpH and Δψ. When ETC function is impaired (such as during lack of terminal electron acceptors or *qox* deletion), or when glucose is in excess, cells favor overflow metabolism (fermentation) to bypass oxidative phosphorylation and produce acidic byproducts. **(B)** Spot titer plates of WT (PY79), Δ*ezrA* (AK140), Δ*qox* (AK430), and Δ*qox* Δ*ezrA* (AK431) strains grown on LB agar alone or supplemented with 2 mM FeCl_3_ or 0.4 mM MnCl_2_. **(C)** Membrane potential (Δψ) of Δ*ezrA* (AK140), Δ*ezrA ezrA^+^* (AK477), Δ*qox* (AK430), Δ*qox* Δ*ezrA* (AK431) was measured using DiOC_2_(3) fluorescence normalized to WT (PY79) within each replicate. Protonophore, CCCP (10 µM), that is known to depolarize cell membrane was used as a positive control. The mean values of three independent biological replicates are shown; error bars indicate standard deviation. Statistical significance was assessed by one-way ANOVA with Tukey’s post-hoc analysis (***, *p* < 0.001; ns, not significant). **(D)** Growth of strains indicated in panel (B) on LB-based phenol red agar, without or with 2 mM Fe^3+^ or 0.4 mM Mn^2+^ supplementation. Phenol red turns yellow upon medium acidification such as when acidic byproducts are made through fermentative metabolism. Colony size and translucency, more obvious in (B), are depicted next to the corresponding strains. Representative plates of three independent biological replicates imaged after 24 h at 37°C are shown.

As ETC function is critical for membrane potential (Δψ) maintenance, we measured Δψ using DiOC_2_(3) assay (Novo *et al*., 1999). As expected, carbonyl cyanide 3-chlorophenylhydrazone (CCCP) protonophore treatment leads to membrane depolarization and served as our control (**Fig. 5C**) (McAuley *et al*., 2018). Using this assay, we see that the Δ*ezrA* strain does not exhibit a reproducible difference in membrane potential when compared to the WT control. In contrast, we see reduced Δψ in the *qox* mutant which is expected as the Qox complex is known to translocate protons (**Fig. 5A**) (Villani *et al*., 1995, Kim *et al*., 2012, Sachla *et al*., 2021). We observe a similar loss in membrane potential in cells lacking *qox* and *ezrA* indicative of compromised ETC function. When respiration is perturbed, cells use fermentative pathways and produce acidic byproducts that lower the media pH (Cruz Ramos *et al*., 2000). To ensure that *qox* deletion does indeed restrict oxidative phosphorylation (**Fig. 5A**), we employed phenol red (PR) as a pH indicator. PR is a standard dye used as a media additive which at neutral pH is reddish pink but turns yellow in acidic pH. In our assay conditions, we see the media pH surrounding WT is in the neutral range whereas the space around Δ*qox* colonies is yellow indicating media acidification (**Fig. 5D**). Furthermore, we see the individual colony diameters of Δ*qox* and Δ*qox* Δ*ezrA* strains are intrinsically small as they are less able to fully utilize respiratory metabolism (Winstedt & von Wachenfeldt, 2000). The difference in colony size between WT and mutants with a *qox* operon deletion is also obvious in **Fig. 5B**. We also note that a Δ*ezrA* mutant forms translucent colonies when grown in the presence of iron and manganese as shown previously (**Figs. 3A** and **5B**). Based on the PR indicator color, we can infer that this mutant is participating in oxidative phosphorylation for energy generation similar to WT. However, when a *qox* deletion is introduced in the Δ*ezrA* background the media pH becomes acidic (PR is yellow) and the colonies appear opaque (**Figs. 5B** and **5D**). To test whether decreased electron flow into the ETC rather than the Qox complex itself allows phenotype reversal, we tested the metal sensitivity of strains lacking *ndh*, which encodes for NADH dehydrogenase (**Fig. 5A**) (Kawai *et al*., 2023, Gaucher *et al*., 2026). As evidenced by the PR indicator, cellular respiration is not fully blocked in this mutant background (**Fig. S4A**), and the colony translucency of the *ezrA ndh* double knockout strain remains uncorrected (**Fig. S4B**). Together, these findings suggest that the Qox complex is the key driver of metal intoxication in cells lacking *ezrA*.

### Promoting overflow metabolism protects Δ*ezrA* cells against metal intoxication

As disrupting ETC function via *qox* deletion alleviates metal-induced toxicity in cells lacking *ezrA* (**Fig. 5B**), we aimed to test this using an independent method. For this, we made use of the knowledge that glucose supplementation favors overflow metabolism (**Fig. 5A**) (Sonenshein, 2007). First, we ensured this was the case in our experimental conditions using the PR plate assay (**Fig. 6A**). For WT and Δ*ezrA* strains the color change from pink to yellow is quite obvious confirming that glucose does promote acidic byproducts production via fermentation. In *B. subtilis*, the phosphoenolpyruvate-dependent phosphotransferase system (PTS) is the major multicomponent glucose importer complex (**Fig. 5A**, bottom panel) (Jahreis *et al*., 2008, Morabbi Heravi & Altenbuchner, 2018). Therefore, as expected, deletion of the *ptsH* gene impaired glucose uptake and colonies remain mostly pink on PR plate even when grown in the presence of excess glucose (**Fig. 6A**). Similarly, we see that the colonies of the double-deletion mutant lacking both *ptsH* and *ezrA* remain pink, indicating the likely utilization of the ETC pathway. As our model suggests ETC-derived ROS as the main source of metal toxicity in the Δ*ezrA* mutant, we hypothesized that glucose supplementation should ameliorate metal intoxication in this strain background. As expected, the typical translucent colonies of cells lacking *ezrA* on iron and manganese plates were more opaque upon glucose addition (**Fig. 6B**). This reversal is not seen when glucose uptake is impaired in Δ*ptsH* Δ*ezrA* double-deletion mutant. Remarkably, glucose supplementation also allows Δ*pfeT* Δ*ezrA* and Δ*mneP* Δ*ponA* mutants to circumvent metal intoxication (**Fig. S3**). We also measured the membrane potential of WT cells in the presence of increasing concentrations of glucose using DiOC_2_(3) assay. This revealed a concentration-dependent decrease in membrane potential indicative of less-active ETC function (**Fig. 6C**). This shows that glucose addition, which facilitates overflow metabolism, alleviates metal-induced toxicity experienced by Δ*ezrA* cells through minimizing the dependency on ROS-prone oxidative phosphorylation (**Fig. 6D**).

**Figure 6:**
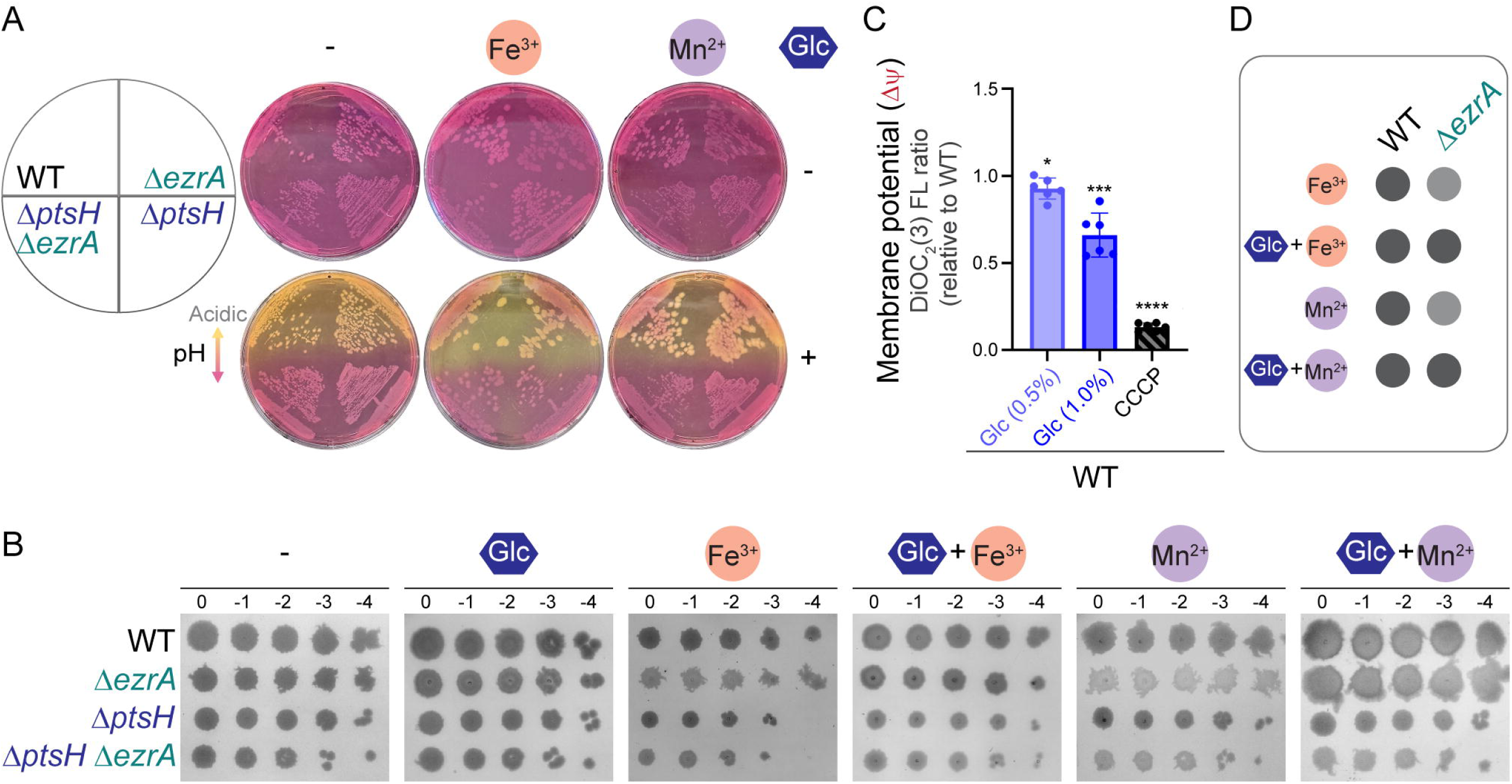
By favoring overflow metabolism, glucose supplementation alleviates metal intoxication in an *ezrA* null mutant. **(A)** Phenol red plates of WT (PY79), Δ*ezrA* (AK140), Δ*ptsH* (AK463), and Δ*ptsH* Δ*ezrA* (AK464) strains grown in the presence of 2 mM FeCl_3_, 0.4 mM MnCl_2_, or neither; without (top) and with (bottom) addition supplementation of 1% glucose (Glc). Yellow color change indicates acidification of the medium. **(B)** Ten-fold serial dilutions of the strains listed in (A) spotted on LB media alone or supplemented with 0.5% glucose, 2 mM FeCl_3_, 0.4 mM MnCl_2_, or in combination as labeled above corresponding plate images. Plates in (A) and (B) were imaged after 16 h incubation at 37 °C and the data shown is representative of three independent experiments. **(C)** Membrane potential (Δψ) of WT (PY79) cells measured using DiOC_2_(3) normalized to untreated cells. Cells were grown in LB with 0.5% or 1.0% glucose; or treated with 10 µM CCCP. Bars represent mean ± standard deviation; n=6. Statistical significance relative to WT was determined by one-sample *t*-test (relative to 1) on log_2_-transformed values; \**p* < 0.05, \*\*\**p* < 0.001, \*\*\*\**p* < 0.0001. **(D)** Schematic summary of colony characteristics for data shown in (B). Increased translucency implies growth impairment due to metal intoxication and this defect is corrected by glucose addition.

Given that glucose and magnesium support the viability of cell wall mutants, does magnesium also inhibit oxidative phosphorylation? We investigated this possibility. In the presence of magnesium, on a PR plate, we did not see medium acidification, but we do observe the colony diameter to be consistently smaller indicative of a slower growth rate (**Fig. S5A**). On the other hand, in non-shaking liquid culture, we notice that the PR color remains in the acidic range (**Fig. S5BC**). Consistent with smaller colonies on plate, we also observe a diminished growth rate with magnesium supplementation in liquid culture (**Fig. S5D**). Upon supplementation of iron to the magnesium containing tubes, the reduced growth rate is corrected, and the PR color change associated with media pH resembled WT. This result further reinforced our model that magnesium restricts growth in stationary culture by inducing iron starvation. Next, we assessed the membrane potential, which was decreased in the presence of magnesium (**Fig. S5E**), but not quite to the same extent as 1% glucose (**Fig. 6C**). Taken together, we believe magnesium hampers ETC pathway. However, as it is not a carbon source like glucose it cannot facilitate overflow metabolism. Consequently, cells grow slowly on magnesium due to iron deficiency and their limited capacity to generate energy.

### Magnesium exposure results in a pronounced reduction in β-lactam susceptibility

Given that magnesium supplementation rescues the viability of cell envelope mutants, we reasoned that this divalent cation may similarly mitigate antibiotic-induced envelope stress and consequently reduce the susceptibility to cell wall-targeting antibiotics. To test this, we used cefepime (Cef), a fourth-generation cephalosporin antibiotic of the β-lactam family (Wynd & Paladino, 1996). β-lactam antibiotics induce cell wall stress by specifically inhibiting the transpeptidase activity of penicillin binding proteins (Mora-Ochomogo & Lohans, 2021). Our results show that WT cells are susceptible to Cef as expected (**Fig. 7A**). Remarkably, this susceptibility is greatly reduced when cells are grown in the presence of magnesium. Conversely, supplementation of iron is not beneficial. As magnesium alleviates iron intoxication (**Fig. 3**), we posited that supplementation of magnesium and iron together should reverse this effect. Indeed, this is what we observe when the cells are grown in the presence of iron and magnesium. As deletion of *qox* also rescues metal-induced toxicity (**Fig. 5B**), we predicted this deletion may also exhibit enhanced resistance to Cef. Again, our result confirms that this is the case. Curiously, however, we see that cells lacking *qox* are less susceptible to Cef only in the absence of magnesium. In fact, Cef susceptibility for this mutant is unchanged with iron supplementation, likely because, without Qox iron-dependent ETC function remains impaired. Additionally, Δ*qox* sensitivity to Cef on magnesium was not alleviated by the addition of iron. We speculate that this is due to a combined effect of restricted ETC function and magnesium-mediated iron starvation. This is particularly detrimental in the presence of Cef. Lastly, as glucose actively facilitates overflow metabolism rather than oxidative phosphorylation and corrects metal intoxication in our mutants (**Fig. 6**), we speculated that it may also render cells less susceptible to Cef. Strikingly, as predicted, this is precisely what we observe. Notably, Δ*qox* cells grow in the presence of glucose as well. We infer this to mean that iron starvation is not induced by glucose addition. These results were also recapitulated in liquid culture (**Fig. S6**). Intriguingly, the protective effect of glucose in liquid culture is not quite as persistent as magnesium (**Fig. S6A**). This is likely because glucose is consumed while magnesium is not. Thus, after glucose exhaustion, cells presumably engage in oxidative phosphorylation and are no longer protected from Cef. Collectively, our results show that Cef sensitivity is plausibly due to dysregulation of ROS-prone ETC pathway. Thus, a reduction in ETC function by glucose via facilitating overflow metabolism or magnesium through inducing iron starvation confers Cef resistance (**Fig. 7B**).

**Figure 7:**
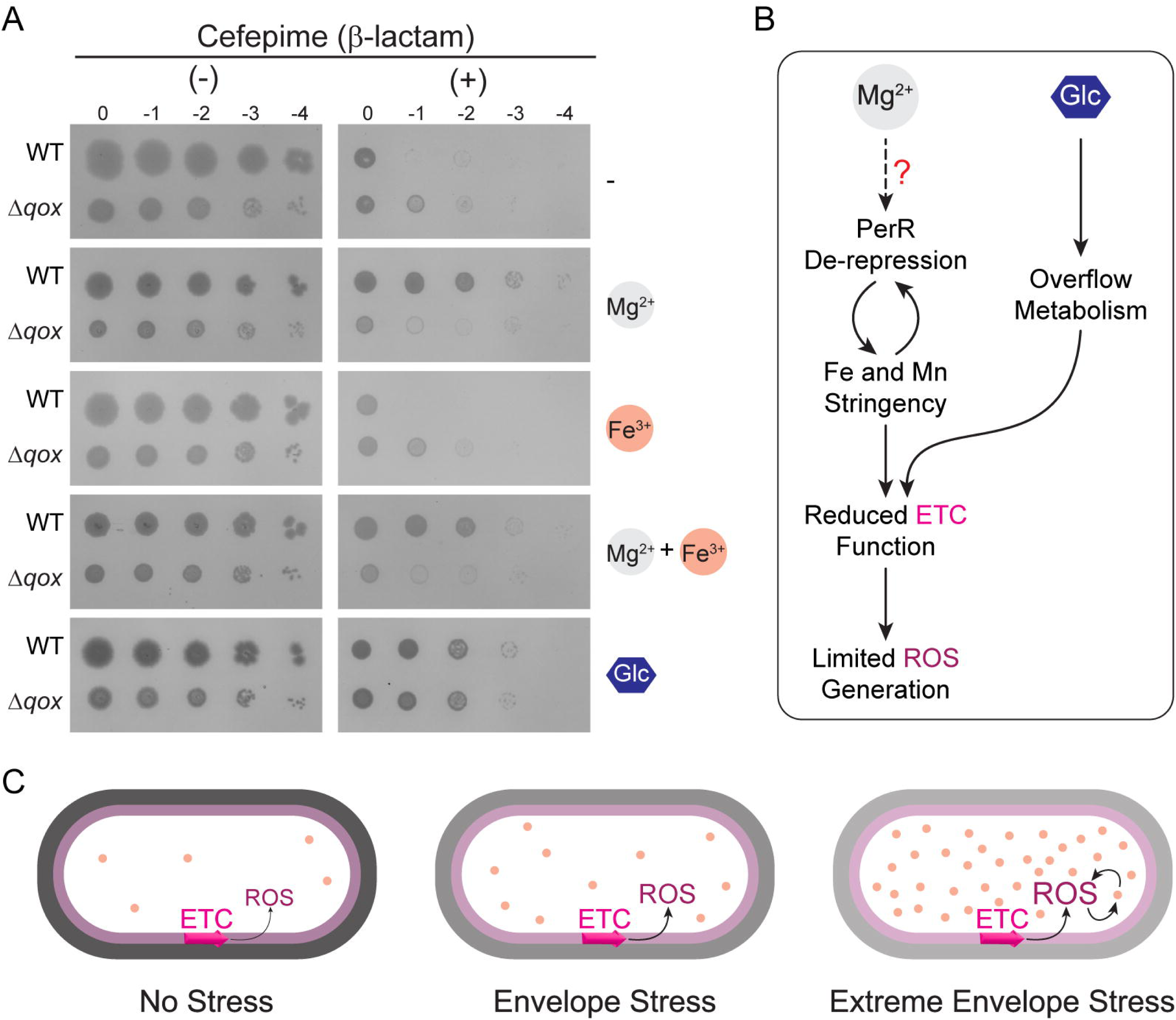
Perturbation of cell wall synthesis impairs ROS-prone ETC pathway the use of which is restricted by glucose or magnesium supplementation. **(A)** WT (PY79) and Δ*qox* (AK430) strains grown without (-) and with (+) the cell wall targeting antibiotic cefepime (0.50 µg/ml) alone (topmost); and in the absence/presence of MgSO_4_ (25 mM), FeCl_3_ (0.2 mM), or both, as well as glucose (1%) are shown. **(B)** Pathway to illustrate the protective effects of magnesium and glucose (Glc). Mg^2+^ likely elicits Fe^2+^ and Mn^2+^ stringency through an unknown mechanism(s), which in turn leads to PerR de-repression responsible for oxidative stress response. PerR de-repression further heightens iron starvation by limiting free iron availability via efflux and/or sequestration. Magnesium-mediated depletion of intracellular iron pool limits the utilization of ETC pathway and minimizes ROS production. Glucose acts independently through metabolic reprogramming towards fermentative metabolism, thereby reducing the dependency on ETC function. **(C)** Model depicting the source of toxicity in cells experiencing envelope stress. In unstressed conditions (left), iron-laden ETC complexes and oxidative stress response pathways function in a synchronized manner to minimize ROS production. Upon experiencing minimal cell wall stress (middle), such as in an *ezrA* null mutant, cells have increased labile iron pool (Fig. 2C), but cells are able to properly activate oxidative stress response pathway (Fig. 1E) to neutralize ROS. However, this ability is compromised when cells encounter extreme cell envelope stress (right), such as in the cases of the *ezrA gpsB* double-deletion mutant or cell wall targeting antibiotic treatment (and also presumably in cells harboring essential cell envelope biogenesis gene disruptions). In this case, our data indicates that reactive radicals stemming from impaired ETC function is further amplified via Fenton chemistry due to increased intracellular labile iron pool. Thus, cell death upon extreme envelope stress is mediated by dysfunctional ROS-prone ETC pathway and impairment of iron homeostasis. We posit that removal of quinol oxidase complex, glucose addition, or magnesium supplementation independently support cell viability by restricting the use of oxidative phosphorylation to limit ROS production and reprogram cells toward fermentative metabolism for energy generation.

## DISCUSSION

Even though the protective effect of magnesium has been known for nearly 50 years (Rogers *et al*., 1976), the precise underlying mechanism has largely remained elusive. In this article, we aimed to address this long-standing puzzle in the field by using a mutant lacking a pair of non-essential genes: *ezrA* and *gpsB*. Although individual deletion of these genes is well tolerated, combinatorial deletion leads to poor growth which is alleviated by magnesium supplementation (Claessen *et al*., 2008). The non-essential nature of these genes allowed us to strategically explore the role of magnesium, which is not straightforward with essential gene disruptions due to their intrinsic requirement for cell survival.

Here, we provided evidence that when both EzrA and GpsB are absent, cells accumulate high levels of ROS and are highly sensitive to oxidative stress (**Fig. 1C**). On a technical note, a recent report encouraged applying an abundance of caution before relating DCF fluorescence with ROS without additional supportive evidence (Korshunov & Imlay, 2025). In our case, the accompanying data showing H_2_O_2_ sensitivity and defective de-repression of the PerR transcription factor strongly suggest ROS involvement in our observed phenotypes. Importantly, we revealed that *katA* expression, which is normally repressed by PerR (**Fig. 1B**), is elevated upon magnesium supplementation for all strains including WT (**Fig. 1E**). Oxidative stress damages iron-sulfur clusters such as those involved in oxidative phosphorylation and results in the release of free iron (Imlay, 2006, Crack *et al*., 2012). Importantly, as elevated levels of iron lead to stronger PerR repression (**Fig. S1D**) (Herbig & Helmann, 2001), we suspected that cells lacking *ezrA* and *gpsB* may also have a high level of intracellular iron. Our results confirmed this prediction (**Fig. 2A**). Additionally, by exploiting the understanding that SN toxicity depends on free iron (**Fig. 2B**), we showed that not only total iron, but also labile iron is elevated in the *ezrA gpsB* double mutant (**Fig. 2C** and **Fig. S7**). This assay also uncovered previously unappreciated roles of EzrA and PBP1 in preventing labile iron accumulation as we noticed heightened sensitivity to SN for the corresponding single-gene deletion strains. Remarkably, the toxic effects of SN in all strains were reduced by magnesium supplementation (**Fig. 2D**). Thus, our results suggest that magnesium reverses SN sensitivity by limiting free iron availability.

If elevated iron levels are responsible for the poor growth in our mutants, then we suspected addition of excess iron, or manganese which also leads to stronger PerR repression as seen for *katA* reporter (**Fig. S1D**) (Herbig & Helmann, 2001), may worsen the growth phenotype. Indeed, addition of iron or manganese to the growth medium leads to stark growth defects in the *ezrA gpsB* double-deletion mutant and delayed growth in an *ezrA* null strain (**Fig. 3A**). Intriguingly, even though cells lacking *ponA* exhibited elevated SN sensitivity, they were almost as resistant as WT to metal intoxication. However, it is to be noted that cells lacking *ponA* constitutively activate an extra-cytoplasmic function sigma factor (SigI) which subsequently upregulates *mreBH* and *lytE* (Patel *et al*., 2020). Thus, the SN and metal sensitivity results cannot be solely attributed to the absence of PBP1. As discussed earlier, if magnesium can restrict free iron availability, we predicted that it may also protect cells lacking *ezrA* individually or in combination with *gpsB* from metal intoxication. Indeed, that is what we observed (**Fig. 3**). This result further strengthens the link between magnesium and metal homeostasis. To further reinforce this possibility, we expanded our study to include metal efflux pump mutants. This set of experiments revealed that the function of PfeT iron exporter becomes critical in cells lacking *ezrA* to alleviate the toxic effects of excess iron (**Fig. 4D**). We saw a substantial growth defect on iron for a *pfeT* mutant with additional deletion of *ponA*. Intriguingly, we found that cells devoid of *ezrA* also rely on PfeT to mitigate manganese toxicity, which was not the case for the *ponA* null mutant (**Fig. 4E**). Furthermore, our results revealed that MneP, the major manganese efflux pump, is important for preventing manganese intoxication especially in cells lacking *ezrA* or *ponA* (**Fig. 4F**). We also see a role for MneS in alleviating manganese intoxication in the absence of GpsB, EzrA, or PBP1 (**Fig. S2**). Astonishingly, all the severe metal toxicity phenotypes are reversed by magnesium supplementation (**Fig. S3** and **Fig. S7**). Collectively, our data demonstrates that limiting the availability of free iron and manganese is the key underlying mechanism behind the protective effects of magnesium (**Fig. 7B**). It is important to note that de-repression of PerR is not the same as *perR* deletion, as the latter results in stunted growth (Faulkner *et al*., 2012).

It was found that the proton-pumping quinol oxidase complex is the sensitive component that leads to manganese intoxication (Sachla *et al*., 2021). Interestingly, it was also demonstrated that either magnesium supplementation or prevention of magnesium export alleviated manganese toxicity (Pi *et al*., 2020, Sachla *et al*., 2024). This motivated us to test whether oxidative phosphorylation (**Fig. 5A**), or more specifically the Qox complex, is also the source of metal-induced toxicity when EzrA is absent. Our experiments established that was indeed the case for both iron and manganese toxicity (**Fig. 5B**). To independently test this finding, we used glucose supplementation which is known to promote overflow metabolism (**Fig. 5A**) (Sonenshein, 2007, Zhuang *et al*., 2011, Jakowec & Finkel, 2025). Consistently, our results revealed that glucose addition restricts the ETC pathway (**Fig. 6AC**). To our delight, glucose supplementation also abolished both iron and manganese sensitivity in cells lacking *ezrA* (**Fig. 6BD**), as well as mutants lacking *ezrA*/*ponA* and metal efflux pumps (**Fig. S3**). Dysregulation of ETC, more specifically the quinol oxidase, is associated with ROS production (Sachla *et al*., 2021). Therefore, the growth rescue of cell wall mutants we observe with *qox* deletion or glucose supplementation can be attributed to limited production of reactive oxygen radicals stemming from the respiratory ETC pathway (**Fig. 7B**). Fascinatingly, we show that both glucose and magnesium are able to independently confer resistance to Cef (**Fig. 7A**). This informed us that ROS generated via oxidative phosphorylation is the primary source of toxicity when cell wall stress is induced by either genetic (Δ*ezrA* Δ*gpsB*) or chemical (cell-envelope targeting antibiotic treatment) means. Although the specific role of ROS in the context of antibiotic-mediated cell death has been a subject of debate (Liu & Imlay, 2013, Dwyer *et al*., 2014, Dwyer *et al*., 2015), our results are in support of this mechanism (Leger *et al*., 2019, Shin *et al*., 2021, Kawai *et al*., 2023, Gray *et al*., 2024).

The role of magnesium in metal homeostasis has been recognized previously (Pi *et al*., 2020, Guo & Herman, 2023, Sachla *et al*., 2024). Interestingly, transcriptomics data of cells treated with magnesium uncovered many genes involved in regulating intracellular metal levels (Guo & Herman, 2023). Notably, among them are genes responsive to iron (Fur regulon) and manganese (MntR regulon) limitation. In conjunction with our results showing that magnesium alleviates the toxic effects of iron and manganese in our mutants (**Fig. 3** and **Fig. S3**), this transcriptomics report further supports our model that magnesium induces iron and manganese deficiency (**Fig. 7B**).

Magnesium is known to affect multiple aspects of cell physiology. Notably, glycerol phosphate polymers (teichoic acids) help assimilate magnesium ions (Heptinstall *et al*., 1970, Lambert *et al*., 1975, Beveridge & Murray, 1980, Thomas & Rice, 2015). Specifically, they bind and concentrate magnesium ions at the cell surface. Consistently, it was shown that magnesium depletion leads to an increase in wall teichoic acid polymer length (Ellwood, 1970, Neuhaus & Baddiley, 2003, Formstone & Errington, 2005). This is believed to increase the binding surface to assimilate magnesium when it is limiting in the environment. Thus, these cell surface polymers are important for maintaining magnesium homeostasis. As such, severe growth defects associated with the deletion of *tagO* which disrupts wall teichoic acid genes are ameliorated with magnesium supplementation (D’Elia *et al*., 2006). Magnesium also affects the cell wall. The availability of cell wall precursors appears to be negatively affected in the presence of excess magnesium (Garrett, 1969, Guo & Herman, 2023, Kawai *et al*., 2023, Dajkovic *et al*., 2017). Similarly, a role for magnesium in peptidoglycan synthesis, hydrolysis and modulation of osmotic stress response has been reported previously (Formstone & Errington, 2005, Chastanet & Carballido-Lopez, 2012, Dajkovic *et al*., 2017, Tesson *et al*., 2022, Wendel *et al*., 2022). Additionally, the effect of magnesium on cell length has also been noted (Webb, 1949, Guo & Herman, 2023). In regard to manganese toxicity mitigation, it was shown that magnesium may act through the shikimate pathway (Sachla *et al*., 2024), which is also responsible for menaquinone production (Shende *et al*., 2024). Specifically, excess magnesium was shown to inhibit the enzymes involved in this pathway (Meneely *et al*., 2016). As menaquinone serves as the main oxygen carrier for aerobic respiration in *B. subtilis* (Yang *et al*., 2020, Chobert *et al*., 2026), it is conceivable that lack of menaquinone may hinder the ETC process. Another possibility is that magnesium may affect the membrane integrity and efficient functioning of ETC components (Reaveley & Rogers, 1969). Intriguingly, we show that membrane potential is negatively impacted by magnesium (and glucose; **Fig. S5E** and **Fig 6C**), which is a key influencer of multiple cellular processes (Benarroch & Asally, 2020). Thus, some of the downstream effects of excess magnesium could be attributed to reduced membrane potential. Regardless, ultimately, we believe the major phenotypic changes are due to iron and manganese starvation especially in cells with compromised cell envelope. However, how magnesium achieves this remains to be elucidated.

In summary, cells under normal growing conditions are well equipped to address the challenges of oxidative phosphorylation. However, cell envelope stress impairs ETC function and leads to ROS production due to increased availability of labile iron such as in the case of the *ezrA* deletion strain (**Fig. 7C**). However, extreme stress such as essential gene disruption or antibiotic treatment, may elevate the intracellular concentration of free reactive iron similar to an *ezrA gpsB* double-deletion strain. The increase in the cytoplasmic labile iron pool and cell wall stress can also be engineered by the combined removal of the major iron efflux pump, PfeT, and EzrA. Such conditions kickstart a destructive positive feedback loop of ROS production through Fenton chemistry. Accordingly, the deleterious effects of Cef, a cell wall targeting antibiotic, are ameliorated by either the deletion of *qox* or by the addition of glucose which limits the use of ROS-prone ETC pathway (**Fig. 7AB**). Similarly, magnesium supplementation also protects, however, in this case the beneficial effect appears to be due to the depletion of free iron.

## MATERIALS AND METHODS

### Strain construction

*B. subtilis* strains utilized in this study are derivatives of PY79 (Youngman *et al*., 1984). Specific details regarding strain construction can be found in the supplemental file. **Table S1** contains all strains and oligonucleotides referenced. The knockout strains were obtained from the Bacillus Genetic Stock Center (Koo *et al*., 2017). All *B. subtilis* chromosomal gene deletions and insertions were confirmed via PCR and other standard techniques.

### Measurement of reactive radical species

Measurement of reactive radical species (RRS) was done using 2’,7’-dichlorofluorescein diacetate (DCFDA) as described previously (Sachla *et al*., 2021). Briefly, single colonies of each strain were selected and grown in 1 ml of lysogeny broth (LB; 10 g NaCl, 10 g tryptone, and 5 g yeast extract per liter) at 37 °C until mid-logarithmic growth phase (OD_600_ ∼0.5). These cultures were back-diluted to OD_600_ of 0.1 in 100 µl in a 96-well plate. Cells were then allowed to grow overnight in a plate reader (BioTek Synergy H1) at 37 °C under shaking conditions and OD_600_ was monitored. DCFDA was added to a final concentration of 1.25 µg/ml and growth was resumed under the same conditions for an additional 6 h. DCFDA fluorescence was measured using excitation and emission wavelength of 498 and 522 nm respectively. DCF-reactive RRS levels were determined by dividing the fluorescence intensity after 6 h by the OD_600_ at the time of DCFDA addition. Values were then normalized to WT levels of fluorescence per OD_600_.

### Hydrogen peroxide disk diffusion assay

Single colonies of each strain were selected and grown in 1 ml of LB at 37 °C until late exponential phase. The OD_600_ of these strains were then measured and each culture was standardized to an OD_600_ of 0.5 in fresh LB. A 100 µl aliquot of each standardized culture was then plated on LB agar and sterile glass beads were used to evenly spread the culture on the surface of the agar plate. Plates were then allowed to dry in a laminar flow hood for 15 minutes. Whatman paper disks were then placed on top of each plate using sterile tweezers. The disks were then laced with 3 µl of H_2_O_2_ (3%) and briefly allowed to dry in the dark. Plates were then incubated in the dark overnight at 37 °C. Zones of inhibition (mm) were then measured the next day using Fiji (Schindelin *et al*., 2012) and analyzed using GraphPad Prism.

### *katA* promoter reporter analysis

Single colonies of the strains to be analyzed were grown shaking in 1 ml of LB at 37 °C until mid-log (OD_600_ ∼0.5). From these cultures, strains were back-diluted to OD_600_ 0.05 into 3 ml of LB amended with additional metals as indicated. These cultures were then grown overnight with shaking at 30 °C. After incubation, 1 ml of each culture was removed and pelleted via centrifugation. The supernatant was removed and cells were washed twice in phosphate buffered saline (PBS). The washed pellet was resuspended in 1x PBS (100 µl) and transferred to a 96-well plate in triplicate. GFP fluorescence was measured in a microplate reader (BioTek Synergy H1) with excitation at 488 and emission at 510 respectively and the observed GFP signal was normalized to the OD_600_ of each sample.

### Spot titer assay

Single colonies of desired strains were grown shaking in 1 ml of LB at 37 °C until mid-log (OD_600_ ∼0.5). Cultures were then back diluted to OD_600_ 0.1 and grown for an additional 60 min. These cultures were then serially diluted (by factors of 10) and plated on LB agar or LB agar amended with various chemicals as needed. Plates were allowed to dry and then were incubated at 37 °C overnight for 24 h or 42 h, or as indicated.

### Phenol red assay

Solid media: Strains were streaked from glycerol stocks directly onto LB agar plates containing 0.002% (v/v) phenol red. Images were taken after 24 h incubation. Liquid media: Single colonies of desired strains were taken and inoculated overnight in 3 ml LB with shaking at 30 °C. The following day a 0.5% (w/v) stock of phenol red was added to a final concentration of 0.002% to LB. To this, the respective overnights of each strain were diluted 1:100. Magnesium and/or iron were added to a final concentration of 25 mM or 200 µM respectively. These cultures were then placed in a 30 °C static incubator and allowed to grow overnight. The following day 1 ml of each culture was taken and centrifuged to obtain cell pellet. The supernatant was removed and stored in fresh tubes and the cell pellet was washed and resuspended in 1X PBS to get an OD_600_ reading that was representative of cell density and not cells with phenol red. OD_600_ was measured for the resuspended cells and each supernatant was measured in tandem looking at the visible spectrum from 400-600 nm. Determination of pH was performed by taking the ratio of yellow (sum of 410-450 nm) to red (sum of 540-580 nm) readings at 10 nm intervals in each tube in comparison to a standard curve.

### Antibiotic assays

Streptonigrin (SN): Single colonies of each strain were selected and grown at 37 °C with shaking in 2 ml LB. Cells were grown until they reached OD_600_ of ∼1 and then back diluted to 0.1 in a 96-well plate. SN (100 µg/ml in DMSO) was then serially diluted in the 96-well plate (by a factor of 3) to create conditions of differing antibiotic concentrations. The 96-well plate was then wrapped in parafilm to prevent evaporation, and the plate was placed in a shaking incubator at 37 °C overnight. Pictures of plates were taken the following day to analyze and document SN sensitivity.

Cefepime (Cef): Single colonies of each strain were inoculated in 2 ml of LB and allowed to grow overnight at 30 °C with shaking. The following day strains were back diluted to 0.1 in 25 ml of LB and then separated into 5 ml aliquots. Magnesium and/or iron were added to final concentrations of 25 mM or 200 µM respectively. Cells in each desired growth condition were then transferred to a 96-well plate to which Cef was added (0.1 mg/mL in water) and diluted down the plate to create differing antibiotic conditions as indicated. The 96-well plate was then placed into a microplate reader (BioTek Synergy H1) and cells were grown at 37 °C shaking for 10 h. Growth dynamics were then analyzed by plotting OD_600_ versus time in GraphPad.

### DiOC_2_(3) Assay

The DiOC_2_(3) assay was performed as described previously (Gentry *et al*., 2010). Briefly, single colonies of each strain were grown in LB to OD_600_ ∼0.5. Cells were then pelleted via centrifugation. The supernatant was removed and the cell pellet was washed with 1 volume DiOC_2_(3) buffer (60 mM Na_2_HPO_4_, 60 mM NaH_2_PO_4_, 130 mM NaCl, 5 mM KCl, 0.5 mM MgCl_2_, 10 mM glucose). Cells were then resuspended in DiOC_2_(3) buffer to an OD_600_ of 3 in 100 µl. DiOC_2_(3) was added to each resuspended pellet to a final concentration of 30 µM and this mixture was allowed to incubate at room temperature in the dark for 5 min. A 20 µl aliquot of this mixture was then diluted in 80 µl of DiOC_2_(3) buffer and placed into a 96-well plate. CCCP (10 µM) was used as a positive control. Imaging was done with an excitation of 450 nm and emission of 490-750 nm (BioTek Synergy H1) after 1 min of incubation in the dark at room temperature. Quantification was performed by taking the sum of the values within the red spectra (600-700 nm) (Gentry *et al*., 2010). A higher value denotes a more robust proton motive force (PMF) while a lower value indicates weaker membrane potential or PMF collapse. Each sample was run in biological and technical triplicate.

### Ferene-S Assay

To measure intracellular iron, individual colonies of each respective strain were grown in LB overnight at 30 °C. Each strain was then diluted 1:100 in 10 ml of fresh LB and allowed to grow to mid-exponential phase OD_600_ ∼0.5. Cells were then pelleted via centrifugation and snap frozen at -80 °C. The day of the assay, respective pellets were thawed at room temperature for 3 min. Pellets were then washed in 1 ml of sterile nanopure PBS. This process was repeated thrice to rinse away residual growth medium. Pellets were then resuspended in 220 µl nanopure PBS containing 1 mg/ml lysozyme and were incubated at 37 °C with gentle shaking for 1 h. Full solution transparency (complete lysis) was ensured before extraction. Extraction was performed via the addition of 25 µl of nanopure HCl (1%) followed by heating to 80 °C for 10 min. Samples were then allowed to cool for 5 min at room temperature. A 200 µl aliquot of each sample was then transferred to fresh 1.5 ml tubes for colorimetric steps. First, 27 µl of ammonium acetate (7.5%) was added to each sample to neutralize excess acid. This was followed by the addition of 27 µl of ascorbic acid (4%) to reduce ferric iron to ferrous iron. Next, 27 µl of SDS (2.5%) was added to assist in the denaturation of any proteins. Lastly, 27 µl of Ferene-S (1.5%) was added to each tube. This mixture was allowed to incubate in the dark at room temperature for 10 min. Then all cell debris were cleared by centrifugation. A 200 µl aliquot of each supernatant was placed in a 96-well plate and absorbance was read at 593 nm, corresponding with the maximum absorbance of the Ferene-S/Fe^2+^ complex (Fritsch *et al*., 2020). Each sample was standardized to the observed OD_600_ of the starting pellet. Standard curves were made in nanopure PBS containing ferrous sulfate to estimate iron content in test samples.

### Statistical Analyses

Statistical analyses were performed using GraphPad Prism v10.4.1 (GraphPad Software, Boston, MA). For comparisons among three or more groups, an ordinary one-way ANOVA followed by Tukey’s multiple comparisons test was used. For data normalized to a single (wild type) reference, values were log_2_-transformed and compared against a hypothetical value of 0 (equivalent to a ratio of 1) using a one-sample *t* test. Comparisons between two samples were assessed by Student’s *t* test. All quantitative experiments were performed with at least three independent biological replicates, each derived from a separate colony and culture; the specific number of replicates for each experiment is indicated in the corresponding figure legend.

## ACKNOWLEDGEMENTS

We are grateful to Dr. John Helmann (Cornell) for strains and for his several publications on metalloregulation which inspired and informed our studies. We thank the members of the Eswara laboratory for comments on this manuscript. This work was funded by the University of South Florida - Center for Antimicrobial Resistance award (P.J.E.) and the National Institutes of Health grant R35GM133617 (P.J.E). The content is solely the responsibility of the authors and does not necessarily represent the official views of the National Institutes of Health.

## AUTHOR CONTRIBUTIONS

Study design (A.K. and P.J.E.), strain construction and data acquisition (A.K. and P.H.), data analysis (A.K. and P.H.), and writing of the manuscript (A.K., P.H., and P.J.E.).

## Notes

### Competing Interest Statement

The authors have declared no competing interest.

