## Supplemental file for "Magnesium induces iron starvation and metabolic rewiring to support the viability of cell envelope mutants and antibiotic-stressed cells"

### SUPPLEMENTAL METHODS

#### Plasmid Construction

To create the plasmid pAK66, we amplified the *ezrA* gene from the *B. subtilis* PY79 chromosomal DNA using primer pairs oAK73/oAK74 (sequence listed below) and cloned it into pDR111 using SalI/NheI cut sites. *Escherichia coli* DH5α was used for cloning purposes.

Oligonucleotide sequences:

oAK73 | 5' *ezrA* primer - SalI

5’-TTATTgtcgacACATAAGGAGGAACTACT*ATGGAGTTTGTCATTGGATTATTAATTGT*-3’

oAK74 | 3' *ezrA* primer - NheI

5’-AATAAgctagc*CTAAGCGGATATGTCAGCTTTGA*-3’

### SUPPLEMENTAL TABLE AND FIGURE LEGENDS

**Supplemental Table 1: Strains used in this study**

**Figure S1: Iron level, streptonigrin sensitivity, and effect of different metal ions on PerR de-repression of *katA*. (A)** Total intracellular iron in WT (PY79) and Δ*gpsB* (GG13) strains, determined by Ferene-S assay. Bars represent mean ± standard deviation (SD) of three independent biological replicates; ns, not significant determined by Student’s *t*-test. **(B)** Streptonigrin (SN) sensitivity in liquid culture. WT (PY79), Δ*gpsB* (GG13), Δ*ezrA* (AK140), and Δ*ponA* (AK286) strains were grown in LB in 96-well plates containing SN at 0, 90, 280, or 830 ng/ml, without or with 25 mM $MgSO_4$. The 96-well plates were imaged after 24 h incubation at 37 °C and data representative of three independent biological replicates is shown. **(C)** Complementation of the Δ*ezrA* SN sensitivity. Top: liquid culture assay as in (B) for WT (PY79), Δ*ezrA* (AK140), and Δ*ezrA* harboring *ezrA* at an ectopic locus (AK477). Bottom: spot titer plates with same strains grown on LB agar with 0 or 50 ng/ml SN, imaged after overnight incubation at 37 °C. **(D)** $P_{katA}$-*gfp* activity of exponential phase WT (AK364) cells grown in LB without or with supplementation of 25 mM $Mg^{2+}$, 0.2 mM $Fe^{3+}$, or 0.2 mM $Mn^{2+}$. GFP fluorescence normalized to $OD_{600}$. Bars represent mean ± SD of three biological replicates; ****$p <$ 0.0001 by one-way ANOVA with Tukey's post-hoc analysis.

**Figure S2: Effects of iron/manganese efflux pump removal on the viability of cells harboring deletions of *gpsB*, *ezrA*, or *ponA*.** Spot titers of strains listed below showing growth at the indicated concentrations of $FeCl_3$ (0, 1, 2, 3, or 4 mM) or $MnCl_2$ (0, 0.4, 1.5, 2, or 2.5 mM) imaged after 42 h at incubation at 37 °C. Images are representative of three independent experiments. **(A)** WT (PY79), Δ*gpsB* (GG13), Δ*pfeT* (AK501), and Δ*pfeT* Δ*gpsB* (AK535) on $Fe^{3+}$ (top) and $Mn^{2+}$ (bottom). **(B)** WT (PY79), Δ*ezrA* (AK140), Δ*ponA* (AK286), Δ*mneP* (AK498), Δ*mneP* Δ*ezrA* (AK513), Δ*mneP* Δ*ponA* (AK532) on $Fe^{3+}$. **(C)** WT (PY79), Δ*gpsB* (GG13), Δ*mneP* (AK498), Δ*mneS* (AK500), Δ*mneP* Δ*gpsB* (AK527), and Δ*mneS* Δ*gpsB* (AK528) on $Fe^{3+}$ (top) and $Mn^{2+}$ (bottom). Red arrowheads (i) mark the $Mn^{2+}$ sensitivity of Δ*gpsB* and Δ*mneS* Δ*gpsB*. **(D)** WT (PY79), Δ*ezrA* (AK140), Δ*ponA* (AK286), Δ*mneS* (AK500), Δ*mneS* Δ*ezrA* (AK511), and Δ*mneS* Δ*ponA* (AK510) grown on LB without (left); or with $Fe^{3+}$ (top right), and $Mn^{2+}$ (bottom right). Red arrowheads mark the $Mn^{2+}$ sensitivity of Δ*mneS* Δ*ponA* (ii) and Δ*mneS* Δ*ezrA* (iii) in comparison to the corresponding individual *ponA* and *ezrA* individual deletion mutants.

**Figure S3: Protective effects of magnesium and glucose in strains lacking genes encoding iron (*pfeT*) or manganese (*mneP*) efflux pump and either *ezrA* or *ponA*.** Spot titer plates of strains indicated below on LB agar alone or with supplementation of 25 mM $MgSO_4$, 1% glucose, 2 mM $FeCl_3$, 0.4 mM (B)/1.5 mM (C and D) $MnCl_2$, or in combination as labeled above the plate pictures. **(A)** WT (PY79), Δ*ezrA* (AK140), Δ*pfeT* (AK501), and Δ*pfeT* Δ*ezrA* (AK504) with $Mg^{2+}$ ± $Fe^{3+}$ (top) and glucose ± $Fe^{3+}$ (bottom). **(B)** As in (A) with $Mg^{2+}$ ± $Mn^{2+}$ and glucose ± $Mn^{2+}$. **(C)** WT (PY79), Δ*ponA* (AK286), Δ*mneP* (AK498), and Δ*mneP* Δ*ponA* (AK532) with $Mg^{2+}$ ± $Mn^{2+}$. **(D)** Strain listed in (C) with glucose ± $Mn^{2+}$. All the plates were imaged after 42 h growth at 37 °C. Representative pictures of three independent biological replicates are shown.

**Figure S4: *ndh* deletion does not protect Δ*ezrA* cells against metal intoxication.** **(A)** Strains of WT (PY79) and oxidative phosphorylation pathway mutants (Fig. 5A), Δ*ndh* (CG204), and Δ*qox* (AK430), grown on LB plate containing phenol red pH indicator. The plate was incubated for 16 h at 37 °C. Yellow color change indicates medium acidification. **(B)** Spot titer plate of WT (PY79), Δ*ezrA* (AK140), Δ*ndh* (CG204), and Δ*ndh* Δ*ezrA* (AK531) strains grown on LB agar alone or with either 2 mM $FeCl_3$ or 0.4 mM $MnCl_2$ imaged after 24 h. Representative plate pictures are shown, n=3.

**Figure S5: Influence of magnesium on membrane potential and overflow metabolism.** **(A)** WT (PY79), Δ*ezrA* (AK140), Δ*qox* (AK430), and Δ*qox* Δ*ezrA* (AK431) strains were grown on phenol red agar plate without (left) or with (right) 25 mM $MgSO_4$. Changes in colony diameter are illustrated next to the corresponding strains. **(B)** Cultures of WT (PY79) strain grown in LB containing phenol red alone or supplemented with 25 mM $MgSO_4$, 2 mM $FeCl_3$, or both. Culture aliquots were photographed after static overnight incubation at 30 °C and representative pictures (n=3) are shown. **(C)** Estimated pH change in culture supernatants of (B). The dashed line indicates the starting pH of the medium (6.5). **(D)** Final $OD_{600}$ value of the cultures in (B). Note: the magnesium-dependent stunted growth phenotype is less apparent with shaking incubation (Fig. S6A). **(E)** Membrane potential (Δψ) of WT (PY79) cells measured using $DiOC_2(3)$ fluorescent probe in untreated cells, upon supplementation of 25 mM $Mg^{2+}$, or treated with 10 µM of CCCP. In (C–E), The means of independent technical replicates (n=3 technical x 3 biological) and corresponding standard deviation (error bars) are shown. Statistical significance was determined by one-way ANOVA with Tukey's post-hoc analysis; **p* < 0.05, ***p* < 0.01, ****p* < 0.001, *****p* < 0.0001; ns, not significant.

**Figure S6: Magnesium supplementation alleviates the toxic effects of cefepime.** Growth curves of WT (PY79) *B. subtilis* grown in LB over a 10 h period with shaking at 37 °C. **(A)** WT cells were grown in the absence (-; left) or presence (+; right) of 0.125 µg/ml cefepime (Cef); and without (left) or with (right) supplementation 25 mM $Mg^{2+}$ or 1% glucose. **(B)** WT strain was grown -/+ 0.125 µg/ml Cef without supplementation or with 25 mM $MgSO_4$, 0.2 mM $FeCl_3$, or both. Representative graphs are shown (n=3).

**Figure S7: Pictographic summary of various phenotypes observed in cell envelope mutants investigated in this study.** Rows indicate the assessed/inferred phenotypes, and columns represent the labeled strain background. N, normal as compared to WT; upward (↑) and downward (↓) arrows indicate increase or decrease relative to WT; double arrows (↑↑ or ↓↓) denote a more pronounced effect; skull symbols denote loss of viability; and asterisks marked not determined due to severe toxicity. The red dashed box, in the Δ*gpsB* Δ*ezrA* background, highlights the elevated levels of ROS, defective *katA* expression (PerR de-repression) and rescue of this defect by $Mg^{2+}$ supplementation which is key to the beneficial effect of magnesium.

### Supplemental Table 1: Strains used in this study

Note: All BKK/BKE non-essential gene knockout strains were obtained from Bacillus Genetic Stock Center (Koo *et al.*, 2017).

| Strain | Genotype | Reference/Notes |
|---|---|---|
| *PY79* | Wild type *B. subtilis* | (Youngman *et al.*, 1984) |
| *GG13* | *gpsB::tet* | (Bhattacharya *et al.*, 2025) |
| *AK140* | *ezrA::cat* | (Bhattacharya *et al.*, 2025) |
| *AK186* | *gpsB::tet ezrA::cat* | AK140 → GG13 |
| *AK286* | *ponA::kan* | BKK22320 → PY79 |
| *AK364* | *amyE::*$P_{katA}$*-gfp spc* | (Hoover *et al.*, 2010) SH517 → PY79 |
| *AK355* | *ezrA::cat amyE::*$P_{katA}$*-gfp spc* | AK364 → AK140 |
| *AK356* | *gpsB::tet amyE::*$P_{katA}$*-gfp spc* | AK364 → GG13 |
| *AK359* | *ezrA::cat gpsB::tet amyE::*$P_{katA}$*-gfp spc* | AK140 → AK356 |
| *AK371* | *ponA::kan amyE::*$P_{katA}$*-gfp spc* | BKK22320 → AK364 |
| *AK430* | *qoxABCD::erm* | (Sachla *et al.*, 2021) HBYL1088 → PY79 |
| *AK431* | *qoxABCD::erm ezrA::cat* | (Sachla *et al.*, 2021) HBYL1088 → AK140 |
| *AK463* | *ptsH::erm* | BKE13900 → PY79 |
| *AK464* | *ptsH::erm ezrA::cat* | BKE13900 → AK140 |
| *AK477* | *ezrA::cat amyE::*$P_{IPTG}$*-ezrA spc* | pAK66 → AK140 |
| *AK476* | *gpsB::tet amyE::*$P_{IPTG}$*-ezrA spc* | pAK66 → GG13 |
| *AK478* | *gpsB::tet ezrA::cat amyE::*$P_{IPTG}$*-ezrA spc* | AK140 → AK476 |
| *AK498* | *mneP::erm* | BKE05470 → PY79 |
| *AK500* | *mneS::erm* | BKE06320 → PY79 |
| *AK501* | *pfeT::erm* | BKE13850 → PY79 |
| *AK504* | *pfeT::erm ezrA::cat* | BKE13850 → AK140 |
| *AK509* | *pfeT::erm ponA::kan* | BKK22320 → AK501 |
| *AK510* | *mneS::erm ponA::kan* | BKK22320 → AK500 |
| *AK511* | *mneS::erm ezrA::cat* | AK140 → AK500 |
| *AK513* | *mneP:: erm ezrA::cat* | BKE05470 → AK140 |
| *AK527* | *gpsB::tet mneP::erm* | BKE05470 → GG13 |
| *AK528* | *gpsB::tet mneS::erm* | BKE06320 → GG13 |
| *CG204* | *ndh::erm* | (Gaucher *et al.*, 2026) |
| *AK531* | *ezrA::cat ndh::erm* | CG204 → AK140 |
| *AK532* | *ponA::kan mneP::erm* | BKE05470 → AK286 |
| *AK535* | *gpsB::tet pfeT::erm* | BKE13850 → GG13 |

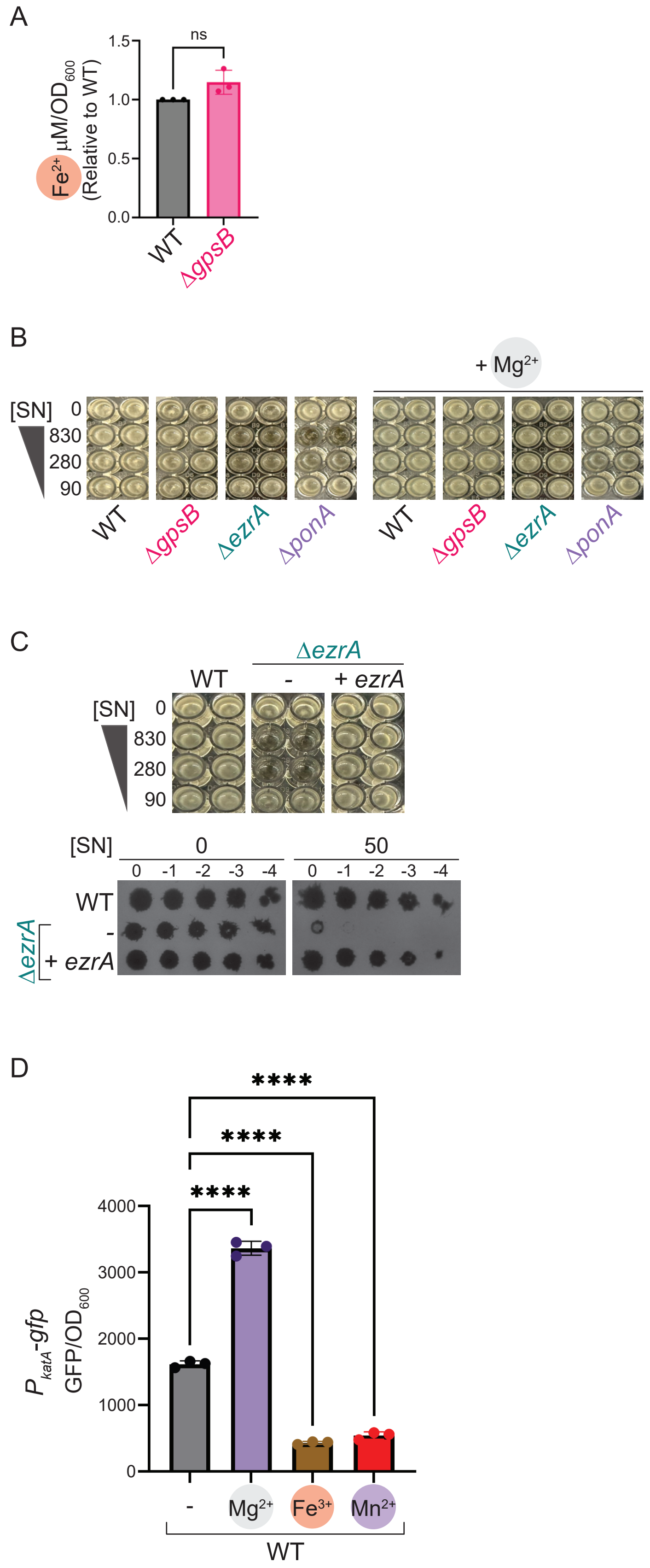
A
ns
Fe2+ µM/OD600 (Relative to WT)
1.5
1.0
0.5
0.0
WT
ΔgpsB
B
+ Mg2+
[SN] 0
830
280
90
WT
ΔgpsB
ΔezrA
ΔponA
C
ΔezrA
WT
-
+ ezrA
[SN] 0
830
280
90
[SN]
0
50
0 -1 -2 -3 -4
WT
-
+ ezrA
D
****
****
****
PkatA-gfp GFP/OD600
4000
3000
2000
1000
0
-
Mg2+
Fe3+
Mn2+
WT

Figure S1

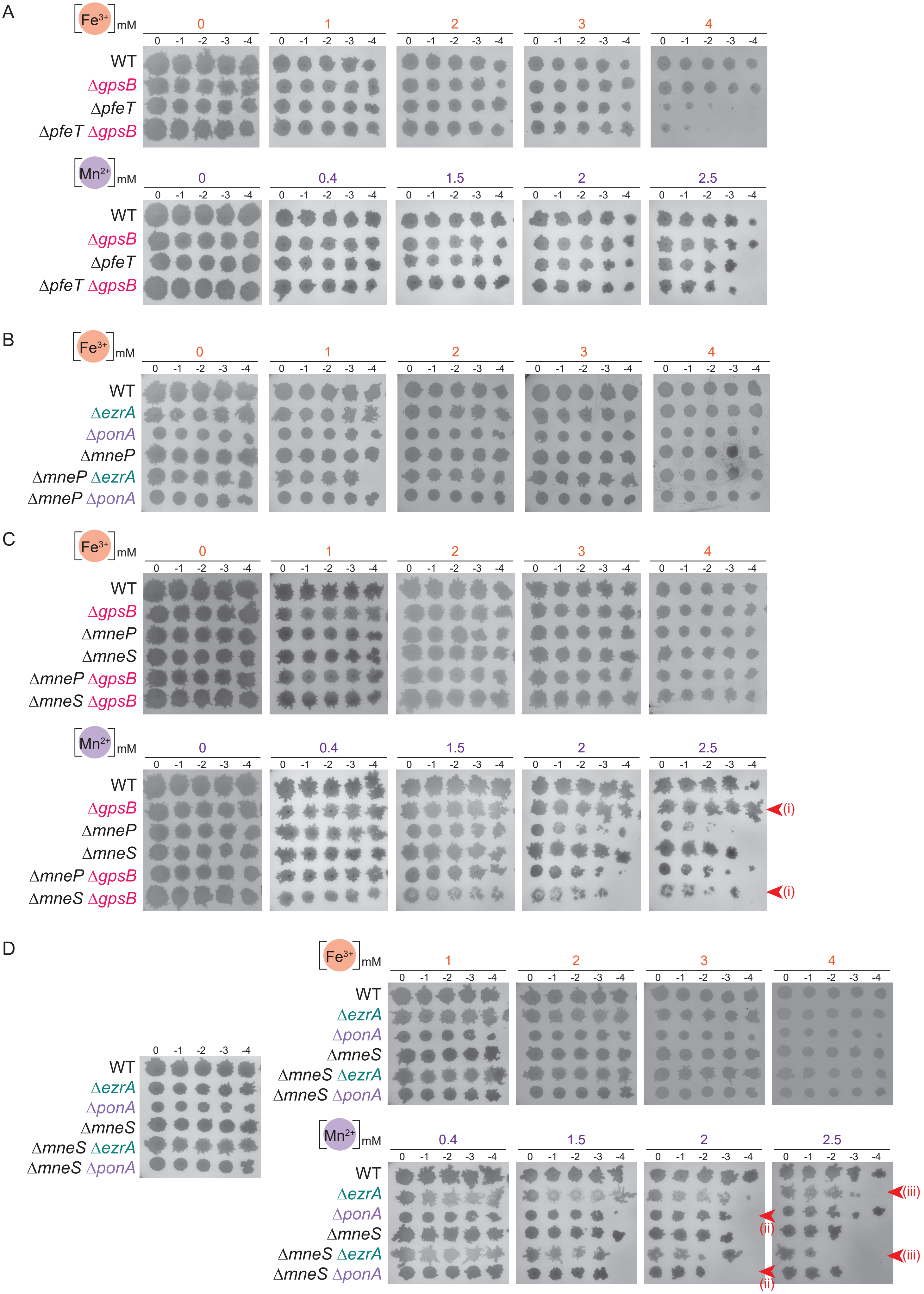

A
[Fe3+]mM
0
1
2
3
4
0 -1 -2 -3 -4
WT
ΔgpsB
ΔpfeT
ΔpfeT ΔgpsB
[Mn2+]mM
0
0.4
1.5
2
2.5
B
ΔezrA
ΔponA
ΔmneP
ΔmneP ΔezrA
ΔmneP ΔponA
C
ΔmneS
ΔmneP ΔgpsB
ΔmneS ΔgpsB
(i)
D
ΔmneS ΔezrA
ΔmneS ΔponA
(ii)
(iii)


Figure S2

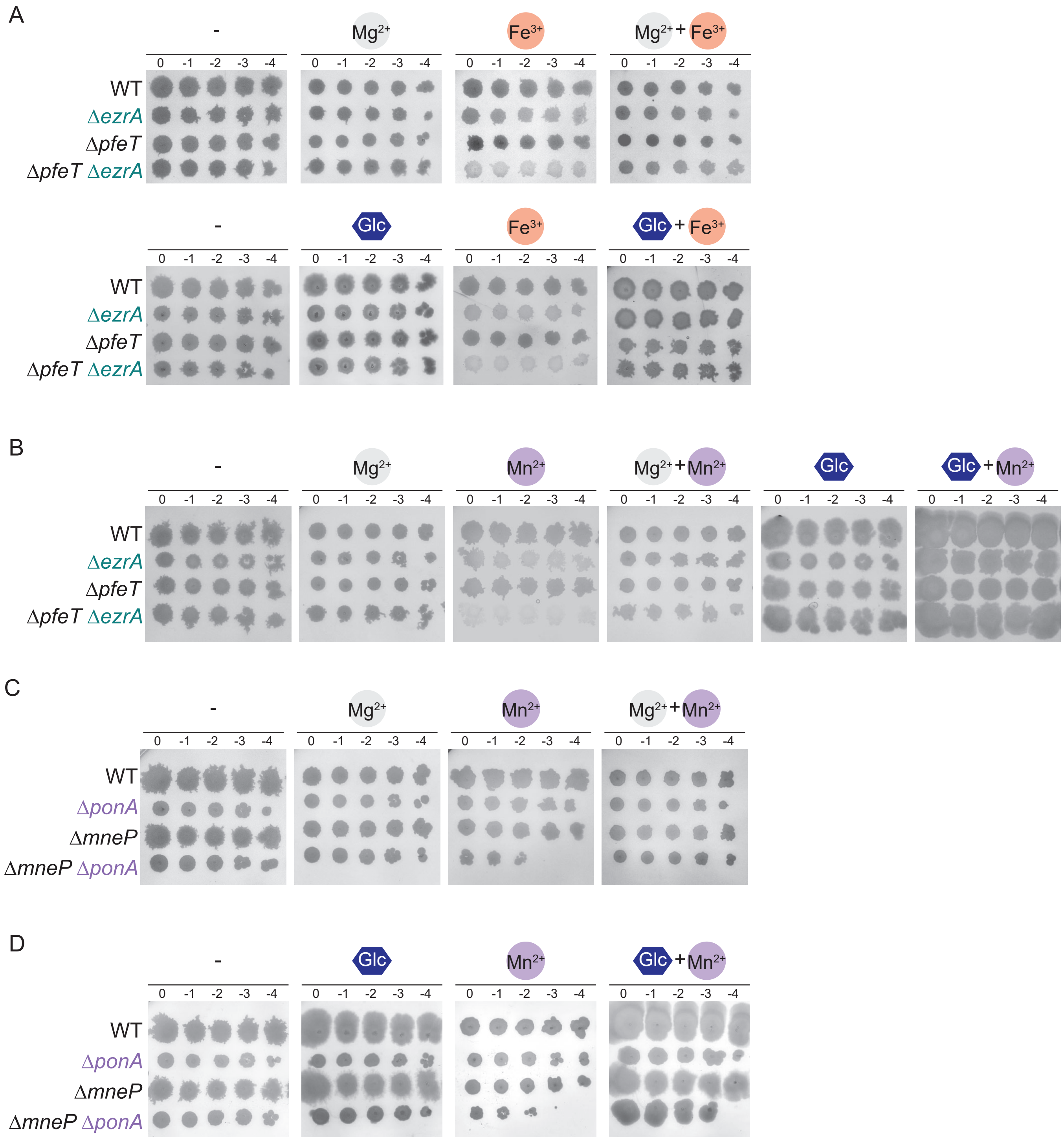
A
-
Mg2+
Fe3+
Mg2+ + Fe3+
0 -1 -2 -3 -4
WT
ΔezrA
ΔpfeT
ΔpfeT ΔezrA
Glc
Glc + Fe3+
B
Mn2+
Mg2+ + Mn2+
Glc + Mn2+
C
ΔponA
ΔmneP
ΔmneP ΔponA
D

Figure S3

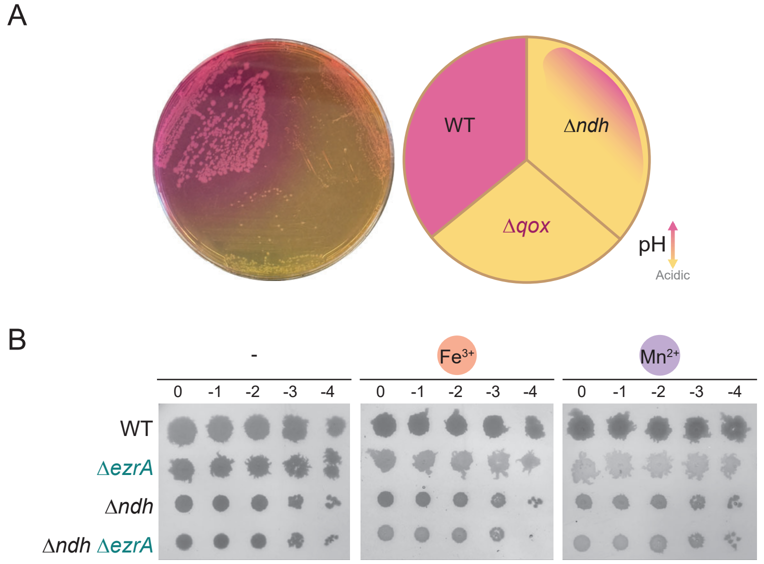
A
WT
Δndh
Δqox
pH
Acidic
B
-
Fe3+
Mn2+
0
-1
-2
-3
-4
WT
ΔezrA
Δndh
Δndh ΔezrA

Figure S4

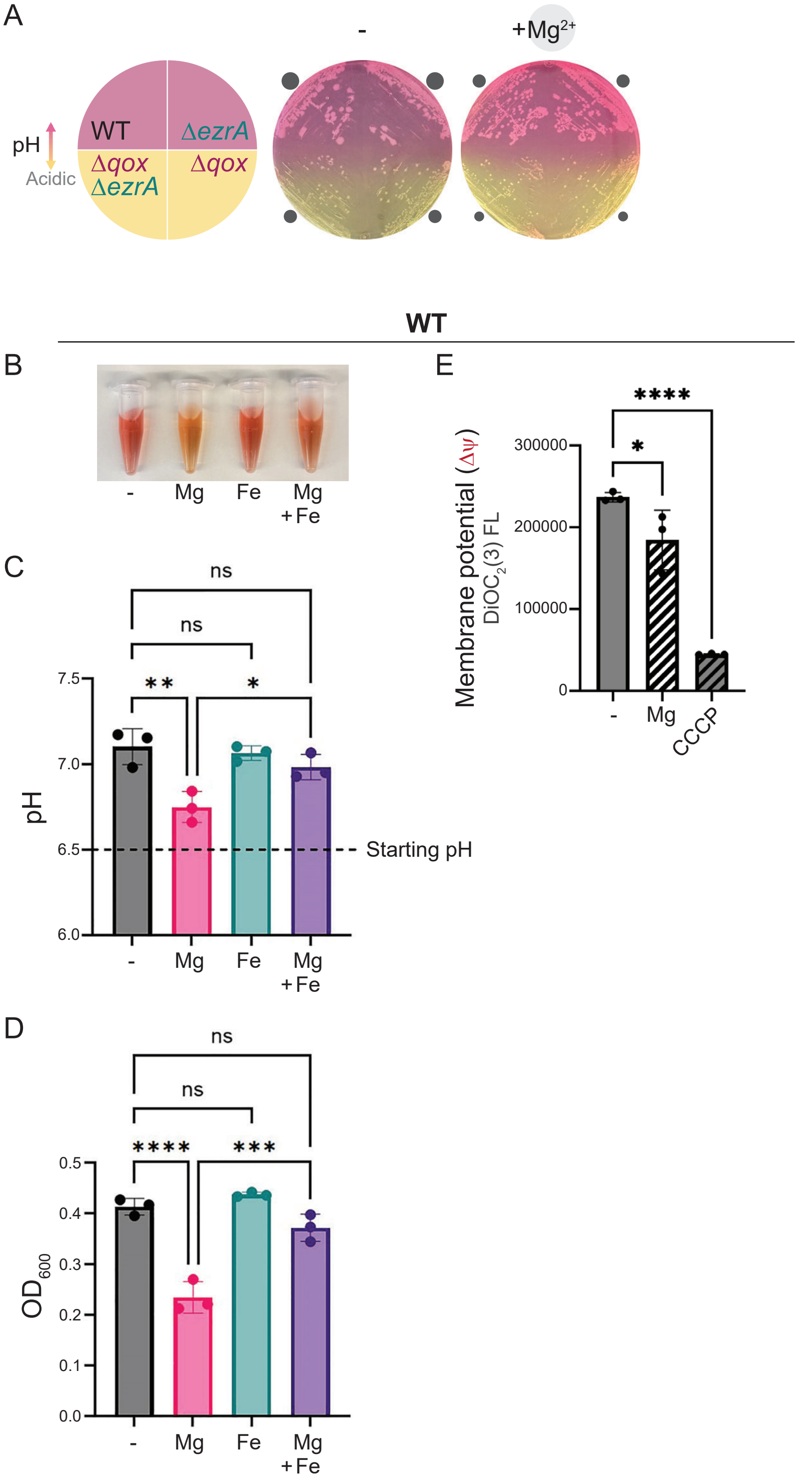
A
-
+Mg2+
pH
Acidic
WT
ΔezrA
Δqox
ΔezrA
Δqox
WT
B
-
Mg
Fe
Mg
+Fe
C
ns
ns
**
*
7.5
7.0
6.5
6.0
pH
Starting pH
D
ns
ns
****
***
0.5
0.4
0.3
0.2
0.1
0.0
OD600
E
****
*
300000
200000
100000
0
Membrane potential (Δψ)
DiOC2(3) FL
CCCP

Figure S5

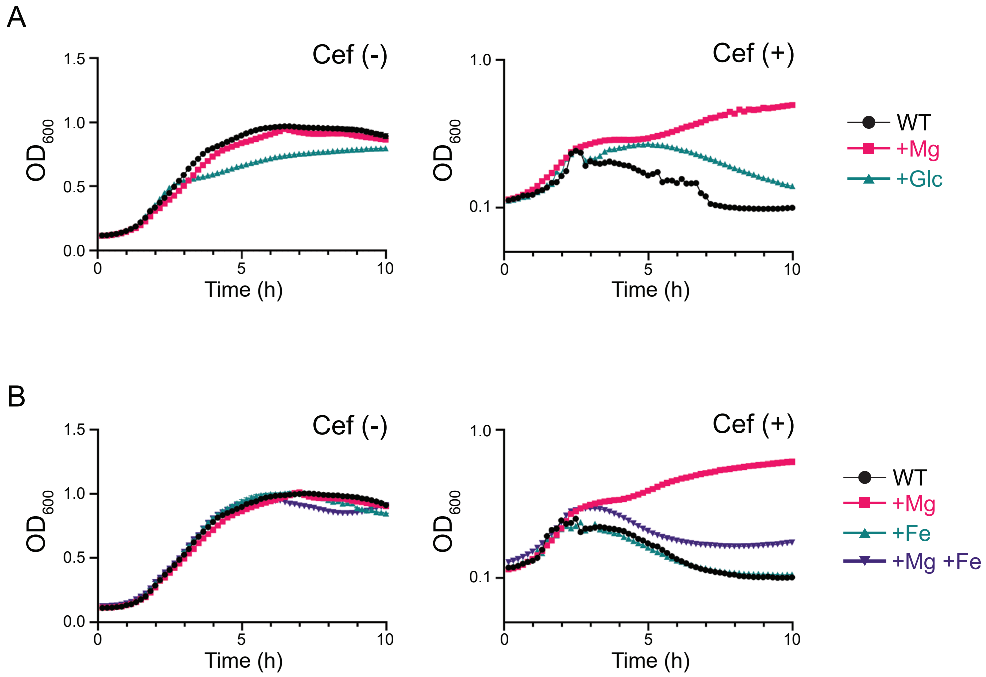
A
Cef (-)
Cef (+)
OD600
Time (h)
WT
+Mg
+Glc
B
Cef (-)
Cef (+)
OD600
Time (h)
WT
+Mg
+Fe
+Mg +Fe

Figure S6

| | | WT | Δ*gpsB* | Δ*ezrA* | Δ*ezrA* Δ*gpsB* | Δ*ponA* |
|---|---|---|---|---|---|---|
| Intracellular labile iron | | N | N | ↑ | ↑↑ | ↑ |
| Streptonigrin toxicity | | N | N | ↑ | ↑↑ | ↑ |
| $H_2O_2$ sensitivity | | N | N | N | ↑↑ | ↑ |
| ROS | | N | N | N | ↑↑ | N |
| *katA* expression | - | N | N | N | ↓↓ | ↓ |
| *katA* expression | $Mg^{2+}$ | ↑ | ↑ | ↑ | ↑↑ | ↑ |
| Sensitivity to | $Fe^{3+}$ | N | N | ↑ | ↑↑ | N |
| Sensitivity to | $Mn^{2+}$ | N | N | ↑ | ↑↑ | N |
| Δ*pfeT* | $Fe^{3+}$ | N | N | ☠ | * | ↑ |
| Δ*pfeT* | $Mn^{2+}$ | N | N | ☠ | * | N |
| Δ*mneP* | $Mn^{2+}$ | N | N | ↑↑ | * | ↑↑ |
| Δ*mneS* | $Mn^{2+}$ | N | ↑ | ↑ | * | ↑ |

Figure S7
